# Commonality of Odorant Receptor Choice Mechanism Revealed by Analysis of a Highly Represented Odorant Receptor Transgene

**DOI:** 10.1101/2022.05.05.489571

**Authors:** Melanie Makhlouf, Charlotte D’Hulst, Masayo Omura, Alessandro Rosa, Raena Mina, Sergio Bernal-Garcia, Eugene Lempert, Luis R. Saraiva, Paul Feinstein

## Abstract

In the mouse, more than 1,100 odorant receptors (ORs) are expressed in a monogenic and monoallelic fashion, referred to as singular gene expression. Using a 21bp singular-choice enhancer (x21), we radically increase representation of olfactory sensory neurons (OSNs) choosing a 5×21 enhanced OR transgene, but not overexpression of its mRNA on a per cell basis. RNA-sequencing and differential expression analysis identified 425 differentially expressed genes (DEGs). ORs make up 86% of all DEGs, of which 325 have decreased representation and 40 have increased representation. Underrepresented ORs include Class I, Class II and TAAR genes and within each of their respective olfactory bulb domains: DI, DII, and DIII (TAAR) we committedly observe multiple homogeneous glomeruli with an OR1A1-identity. The underrepresentation of endogenous, class-specific ORs across evolutionarily distinct cell types in favor of the expression of the 5×21-OR1A1 transgene argues that a common mechanism of singular gene choice is present for all OR-expressing OSNs.

## Introduction

The mouse main olfactory epithelium (MOE) is populated by ten million olfactory sensory neurons (OSNs), expressing a total of approximately 1,100 odorant receptor (OR) genes (Ibarra-Soria et al., 2014; Zhang et al., 2004). Each mature OSN expresses predominantly one allele of a single OR gene, thus defining the identity of the thousands of different OSN subtypes (Tan et al., 2015; Tsukahara et al., 2021), with each type projecting axons that coalesce into a homogeneous glomerulus in the olfactory bulb (OB). As such, it is said that the mechanism of singular gene choice governs the expression of one allele out of a repertoire of over 2,200 OR alleles. While the mechanism of OR expression through singular gene choice remains poorly understood, two features codify this phenomenon; exclusive expression in OSNs and coalescence of their axons to form homogeneous glomeruli (Feinstein et al., 2004; Feinstein and Mombaerts, 2004). Here we focus on the expression profile of an OR transgene expressed broadly among 13% of all OSNs (D’Hulst et al., 2016).

OR genes (*Olfrs*) and TAAR genes have three defining characteristics: i) they are G-protein coupled receptors (GPCRs) with seven transmembrane domains, ii) they display stable singular gene expression in OSNs, and iii) they promote the coalescence of like axons into homogeneous glomeruli in the main OB (Feinstein and Mombaerts, 2004).

When OR genes were first cloned, it became apparent that they fell into two phylogenetic categories, designated Class I and Class II ORs (Zhang and Firestein, 2002; Zozulya et al., 2001). Experimental evidence showed that a non-OR 7-transmembrane GPCR gene, β2adrenergic receptor (β2AR), when expressed in OSNs from the Olfr151 [M71] locus (β2AR→Olfr151-IRES-tauLacZ) could lock in singular gene choice and define axonal identity resulting in the generation of functional homogeneous glomeruli in the OB (Feinstein et al., 2004; Omura et al., 2014). A third class of 15 OR-like genes were identified in mouse that had homology to the trace amine-associated receptor 1 (Taar1) (Liberles and Buck, 2006). Despite TAARs having little sequence homology to Class I and Class II ORs, they follow the aforementioned characteristics of ORs.

Singular expression for a given OR gene is normally restricted within regions of the MOE. This restriction is referred to as stochastic gene expression within a zone. There has been much debate as to the number of zones within the MOE. Early studies using in situ hybridization techniques identified four zones and a “patch”(Ressler et al., 1993; Strotmann et al., 1992; Vassar et al., 1993). In an experiment using 100 OR FISH probes, it was deemed that all ORs can be ascribed as having their own expression pattern along the dorsomedial/ventrolateral axis (Miyamichi et al., 2005). A study combining regional dissections of random small pieces of the MOE coupled with RNA-sequencing (RNA-seq) mapped 1,033 mouse ORs and seven TAARs to 39 distinct dorsal to ventral (DV) regions (Zone Indexes 1.05 to 5.0) and an “unusual location” (patch zone) (Tan and Xie, 2018). Similarly, another recent study using a spatial transcriptomics approach created a genome-wide 3D atlas of the mouse whole olfactory mucosa (WOM) and found that ORs are distributed in a continuous fashion over five epithelial regions (Ruiz Tejada Segura et al., 2022). In contrast, another study combining in situ hybridization of approximately 70 OR genes with wholemount 3D reconstructions, suggested that the minimum number of OR zones is nine (Zapiec and Mombaerts, 2020). Of all the methods of delineating zones to date, the indices calculated from RNA-seq performed on pieces of MOE collected along the dorsoventral axis represents the least biased approach for landmarking the topological limits of particular OR gene expression (Tan and Xie, 2018). The addition of data from lateral-medial and anterior-posterior axes may help further refine these indices to produce a granular map of OR expression in the MOE.

All previous studies have used the rationale of topological expression patterns of an OR for the determination of zones, however, none have provided a mechanism for how the zone that a cell inhabits influences OR choice and expression. One biological rationale for zones could be to mechanistically reduce the complexity of the problem of singular gene choice. The definition of zone was expanded by showing that the same OR gene, broadly expressed by OSNs in adjacent “zones” had different axonal identities and formed multiple homogeneous glomeruli in the OB (Feinstein and Mombaerts, 2004). These data argued that broader expression of an OR across the MOE revealed cell-type identities. A subsequent study definitively showed that Class I OR genes, despite being expressed in overlapping regions of the MOE with Class II OR genes within the dorsal MOE, projected to a distinct Class I “domain” (DI) of the OB (Bozza et al., 2009). Additionally, when a deletion Class I OR allele undergoes second choice, the subsequent choice is restricted to a Class I OR gene, which also projects to this same Class I region in the OB. These OR deletion alleles seem to reflect cell-type identities; that is a Class I OR deletion allele coexpresses with another intact Class I OR (Bozza et al., 2009). It was then established that TAAR genes are also dorsally restricted, overlap in expression with Class I and Class II ORs in the MOE, and project to a third distinct “TAAR” region (DIII) in the dorsal bulb (Pacifico et al., 2012). As with the deletion Class I OR allele, OSNs making the first choice of a deletion TAAR allele (ΔTAAR) undergo second choice which is restricted to intact TAAR genes, and then go on to project axons to the same TAAR domain in the OB (Pacifico et al., 2012). Therefore, Class I ORs, Class II ORs, and TAARs all reside in overlapping regions of the dorsal MOE, show class restriction in singular gene choice, and group their axonal projections to different domains in the OB. Of consequence, a Class II OR coding sequence expressed by a Class I OR promoter projects axons to and forms glomeruli in the Class I domain of the OB (Bozza et al., 2009), further arguing that it is the transcriptional regulatory regions of Class I, Class II, and TAAR OR genes that dictate cell-type restriction, not the receptor coding sequence (Feinstein et al., 2004). Thus zones, or regions of OR expression, as originally described, are not a defining characteristic of class restriction of OR gene expression as multiple cell-types coexist within the same topological location in the MOE.

As outlined above, cell-type restriction of an OR gene can be defined in two ways. First, by the probability that a given OR gene can be expressed in an OSN, visualized by the repertoire of ORs expressed in OR deletions as a second choice. Second, by the formation of segregating glomeruli in the OB. As an example, Olfr151 is normally expressed in Class II OCAM-dorsal cells and does not project within the Class I domain of the OB. We previously showed that when a high-probability element (4×21-Olfr151α), derived from a four-times multimerization of the 21bp core homeodomain site (HD) within the H-enhancer element (x21), drives expression of a minigene, there is a substantial increase in the probability of choice of the transgene, but Class II cell-type restriction is maintained (D’Hulst et al., 2016). Unlike the canonical Greek Island motif (Monahan et al., 2017), where an HD site is found within 3bps of an O/E-like site, our x21 element does not contain a proximal O/E-site. We went on to show that when we increase the probability of choice to 13% of the MOE with our hemizygous, H-derived, 5×21-OR1A1 minigene line, we found a number of distinct glomeruli that appear to innervate both the Class I and Class II regions of the OB and line up in a row (D’Hulst et al., 2016), thereby breaking cell-type restriction. At the time, we could not rule out the possibility that the breaking of cell-type restriction was the result of the use of a human OR coding sequence in our high probability transgene. We subsequently showed that a 5×21-transgene with no OR sequence also sends axonal projections to both Class I and II regions of the OB, consistent with the 5×21 enhancer element being a substrate for breaking cell-type restriction (Shah et al., 2021).

If a Class I choice element could be added to the Class II Olfr151 minigene or if the Olfr151 promoter used were to somehow gain a Class I probability of choice, then one would expect to see multiple distinct glomeruli in the OB; ones that were exclusively co-innervated with the endogenous Olfr151 Class II alleles in the Class II regions of the bulb and one in the Class I region of the OB, exclusively co-innervated by axons expressing the Olfr151 minigene. Here we show that using a germline-expressed transgene identical to 4×21-Olfr151α (4×21-Olfr151β mouse line) and randomly integrated in the X-Chromosome, cell-type restriction is broken, and we see co-innervation of endogenous Olfr151 axons with transgene axons in the Class II glomeruli and the formation of a transgenic Class I glomerulus, similar to what we observed with 5×21-OR1A1.

It is important to note that **high probability of choice** is defined as the significant **overrepresentation** of the number of cells within the MOE monoallelically expressing a particular OR gene, and **not** the result of overexpression in the classical sense of gene amplification within a given OSN or coexpression with endogenous ORs (D’Hulst et al., 2016). RNA levels of the OR gene transcribed within high-probability choosing cells are equivalent to those that chose to express the endogenous OR allele (D’Hulst et al., 2016). This is confirmed by the maintenance of axonal identity and the coalescence of axons from OSNs within the same cell type expressing either the endogenous or high-probability driven alleles projecting to the same homogeneous glomerulus in the OB and also suggests that OR protein levels are equivalent in both populations of cells allowing for concomitant axonal targeting (Feinstein et al., 2004; Feinstein and Mombaerts, 2004). Therefore, Class II OSNs expressing the x21-Olfr151 transgenes have the same identity as Class II OSNs that express the endogenous Olfr151 alleles. As we show here with our 4×21-Olfr151β and 5×21-OR1A1 lines, high probability of choice elements are also capable of breaking cell-type restriction and significantly increase their representation within the MOE by coopting choice in other OSN cell types. Prior models of singular gene expression posited that class-specific restriction arose from unique mechanisms for OR choice (Bozza et al., 2009; Enomoto et al., 2019; Pacifico et al., 2012), however, the data we present herein suggests that there is in fact a common mechanism of OR choice which underlies singular gene expression for all of the three main OR classes.

Here, we performed RNA-seq on the MOE of hemizygous 5×21-OR1A1 and wild-type mice and identified the differentially expressed genes (DEGs) between both strains. Among genes that are significantly underrepresented in 5×21-OR1A1 vs. wild-type we found Class I ORs, Class II ORs, and TAARs, arguing that the 5×21-OR1A1 minigene has gained probability of choice in those OSN cell types and has broken the regime of cell-type restriction based on OR class. The observed reduction of Class I OR expression and increased OR1A1 representation in the dorsal epithelium is not simply an expansion of Class II OSNs in the MOE as no gains in Class II expression were observed in mice with reduced Class I OR repertoires (Cichy et al., 2019; Iwata et al., 2017). Our data argue for four important findings concerning OSN cell types: i) cell types create distinct probabilities of choice for a given OR gene; ii) cell types dictate a codifying axonal-identity property whereby an OR that gains an additional cell-type probability leads to the formation of unique glomeruli; iii) the total number of OSN cell types can be ascertained by the expression of a gene that has broken all cell-type restriction and is represented broadly across the entire MOE, leading to the innervation of multiple unique glomeruli; and iv) that there are multiple subclasses of the Class II cell type, suggesting that there are likely multiple TAAR and Class I cell types as well. Finally, our data argue that there is a commonality in the mechanism of singular gene choice across all chemosensory GPCR-expressing cell types in the MOE. There are, however, cell-type specific modulators that regulate probabilities of choice so that only a subset of ORs are selectable at a high enough probability such that there are sufficient numbers of mature OSNs with the correct identity projecting axons to form stable glomeruli. We hypothesize that our high-probability element transgenes can overcome the cell-type specific modulators, possibly as a result of enhanced transcriptional accessibility through the cooperativity of multimerized regulatory elements, access the common mechanism of choice for all OR and TAAR-expressing cell types, and break the cell-type restriction regime, allowing them to be expressed broadly in the MOE. These high-probability choice elements will be an important tool to probe the mechanism of olfactory gene choice and allow us to determine the different OSN cell types in the MOE.

## Results

### Expression of odorant receptor transgenes

#### 5×21-OR1A1 transgene

Our previously published 5×21-OR1A1→Olfr151-IRES-tauCherry (5×21-OR1A1) hemizygous transgenic line with a coding sequence swap of OR1A1 for Olfr151 stood out from the Olfr151 lines (D’Hulst et al., 2016) in two important ways that may have been correlated. First was the broad distribution of OR1A1 expression in the MOE with approximately 13% of OSNs expressing the transgene in a monogenic manner; that is one million OSNs in the MOE exclusively expressing a single OR transgene. Second was the formation of multiple homogeneous glomeruli within the same domain of the OB and appeared to enervate multiple class domains of the dorsal bulb as defined in prior studies (Pacifico et al., 2012). During early postnatal development, distribution of OR1A1 expression appears evenly dispersed (PD15, **Figure 1A**), but by PD30, the majority OR1A1-expressing OSNs are found in the dorsal MOE (**Figure 1A inset**). Crosses to a TAAR deletion allele (ΔTAAR4-YFP) line that labels all dorsal TAAR glomeruli (Pacifico et al., 2012) reveal yellow fluorescent neurons confined to areas of dense OR1A1 expression in the dorsal MOE (**Figure 1B**). Broad distribution of OR1A1 expression in the MOE correlated with many medial glomeruli on the posterior bulb resembling an island chain (**Figure 1B**, asterisks). Of note, the dorsally-focused ΔTAAR4-YFP axons (yellow) bisect this grouping of OR1A1 glomeruli (red) with two OR1A1 glomeruli dorsal of and at least seven OR1A1 glomeruli ventral to the TAAR fibers. Also, the multiple OR1A1 glomeruli appear regularly spaced and positioned equally posterior. Like the medial projections, the lateral projections on the dorsal bulb are distributed in an island chain pattern as well (**Figure 1C**). We made a triple transgenic reporter line by crossing 5×21-OR1A1 (red-OR1A1 axons) to both P-LacZ (Bozza et al., 2009) (blue-Class II (DII) axons) and ΔTAAR4-YFP (green-TAAR (DIII) axons). This line revealed additional red axons that are not Class II axons and are dorsal to TAAR axons (**Figure 1D-G**), suggesting a third cell type expressing the OR1A1 transgene.

**Figure 1.**
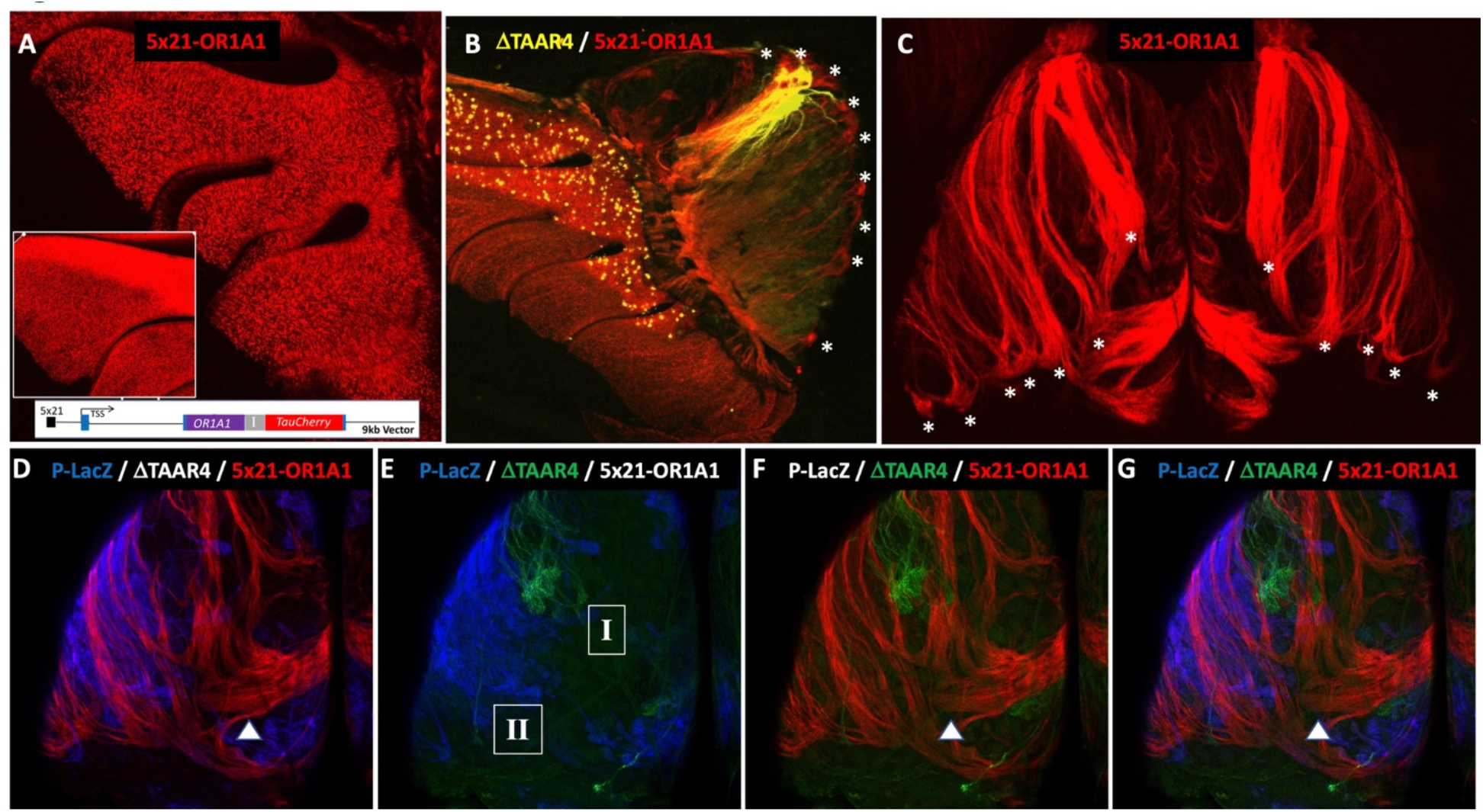
Epithelial expression and axonal projection in 5×21-OR1A1-IRES-TauCherry (5×21-OR1A1) transgenic mice. (A) Confocal medial wholemount view reveals distributed red fluorescence in epithelial turbinates during early development, PD15, which becomes significantly denser in the dorsal MOE by PD30 (inset). (B) Confocal medial wholemount view (PD30) for 5×21-OR1A1 crossed to ΔTAAR4-YFP (TAAR axons (DIII)-yellow) reveals dense red fluorescence in dorsal MOE and projecting axons to multiple distinct medial glomeruli on the posterior bulb having an island chain pattern marked with asterisks (*). TAAR axons (yellow) separate the most dorsal red glomeruli from ventral glomeruli. (C) Confocal dorsal wholemount view of OBs for 5×21-OR1A1 transgene reveals projecting axons to multiple distinct lateral glomeruli on the posterior dorsal bulb (*) similar to the medial projections seen in 1B. (D-G) Channel-merged confocal dorsal wholemount view of the same OB for 5×21-OR1A1 (OR1A1 axons-red) crossed with P-LacZ (Class II axons (DII)-blue) and TAAR4-YFP (TAAR axons (DIII)-green). (D) The most dorsal red glomerulus (arrowhead) does not reside in the Class II-blue region. (E) The TAAR axons-green do not reside in the Class II-blue region (II). The area identified by I is the putative Class I domain of the OB (DI). (F-G) A large red glomerulus (white arrowhead) resides dorsal to Class II and TAAR axons, suggesting a Class I origin.

#### 4×21-Olfr151β transgene

We previously dissected the elements of OR gene expression through the generation and characterization of OR minigenes for Olfr16 [MOR23] and Olfr151 [M71] (Vassalli et al., 2002). Dissection of the control regions for six additional OR promoters led to the identification of a 21bp sequence that, when multimerized, dramatically increased the probability of representation for a given Olfr151 transgene in the MOE (D’Hulst et al., 2016). These 4×21-Olfr151-IRES-tauCherry alpha (4×21-Olfr151α) and 5×21-Olfr151-IRES-tauCherry alpha (5×21-Olfr151α) lines exhibited a 40-fold increase in OSNs expressing Olfr151, while recapitulating the expression pattern of Olfr151-IRES-tauGFP knock-in lines (D’Hulst et al., 2016). The transgene-expressing OSNs maintained their Class II, Olfr151 identity, even with the substantial increase of cell representation in the MOE; their axons coalesced with Olfr151-IRES-tauGFP axons within the same glomeruli (D’Hulst et al., 2016). We observed the formation of multiple Olfr151 (Cherry) homogeneous glomeruli in 5×21-Olfr151α animals, however, due to the fact that the observations were made through founder analysis, we could not rule out that the phenotype was the result of the lack of imprinting in animals derived by transgenic pronuclear injection. Here we describe an additional germline Olfr151 transgenic line, 4×21-Olfr151-IRES-tauCherry beta (4×21-Olfr151β), identical in sequence to 4×21-Olfr151α and randomly integrated into the X-Chromosome, that exhibits an increase in OSNs expressing the transgene compared to the alpha line, but remain dorsally restricted. In these mice, transgene-expressing OSNs project to multiple defined glomeruli (**Figure 2A**) and appear to express the transgene in a monogenic fashion; when crossed to an Olfr151-IRES-tauGFP knock-in, OSNs express either the transgene or the Olfr151 endogenous allele.

**Figure 2.**
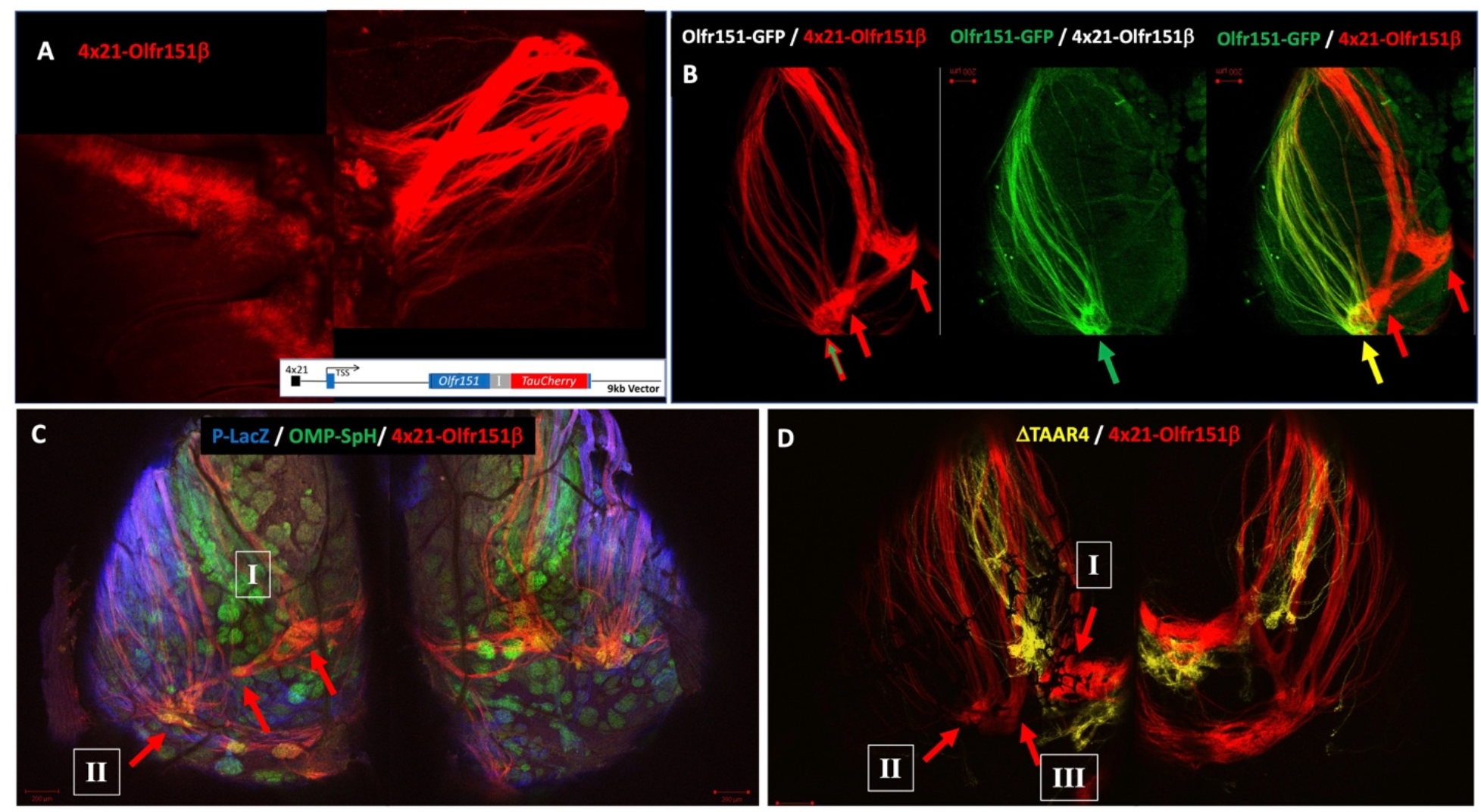
Epithelial expression and axonal projection in 4×21-Olfr151-IRES-TauCherry-β (4×21-Olfr151β) transgenic mice. (A) Confocal medial wholemount view reveals a dense swath of red fluorescence in the dorsal MOE and medially projecting axons to multiple glomeruli on the posterior dorsal bulb. (B) Confocal dorsal wholemount view of OB for 4×21-Olfr151β crossed with Olfr151-IRES-TauGFP knock-in reveals lateral projections of green fluorescent axons to only one of the three red glomeruli on the dorsal surface. (C) Confocal dorsal wholemount view of OBs for 4×21-Olfr151β crossed with P-lacZ (Class II axons (DII)-blue) and OMP-SpH (green glomeruli). Only one of the three red glomeruli resides in the Class II domain (II) of the dorsal bulbs. (D) Confocal dorsal wholemount view of OBs for 4×21-Olfr151β crossed to βTAAR4-YFP (TAAR axons-yellow). The most lateral of the three red glomeruli coincides with the Class II domain (II). A second red glomerulus (III) resides within the TAAR axon projections, which are not of Class II origin and are more dorsally focused. A third large red glomerulus is more medially positioned than the TAAR axons, suggesting a Class I origin (I).

One unusual property of this line was the consistent formation of three dorsal glomeruli, innervated by axons from OSNs located in the dorsal MOE. This suggested that the transgene may be expressed across the three known dorsal OSN subtypes with axonal projections to the DI, DII, and DIII (TAAR) domains of the OB (Pacifico et al., 2012). To test this idea, we asked whether the lateral-most glomeruli were formed by endogenous Class II OSNs, the OSN subtype that normally expresses Olfr151. We crossed the 4×21-Olfr151β line to the Olfr151-IRES-tauGFP knock-in. In these mice, we observed co-innervation of red and green axons only in the most lateral of the dorsal glomeruli (**Figure 2B**), which corresponds to the DII domain. To further test whether these lateral glomeruli were part of the DII domain, we crossed the 4×21-Olfr151β line to a P-LacZ transgene line which co-expresses the beta-galactosidase enzyme broadly in 10% of all Class II OR expressing OSNs and labels DII axons (Bozza et al., 2009). We observed that two Olfr151β-enervated glomeruli reside outside the Class II domain of the dorsal bulb (**Figure 2C**). Finally, to determine if one set of glomeruli correspond to TAARs, we crossed the 4×21-Olfr151β line to ΔTAAR4-YFP and observed that TAAR axons lay next to a red Olfr151β-enervated glomerulus centered between the DI and DII domains (**Figure 2D**).

These data suggest that the 4×21-Olfr151β expression pattern is consistent with it being expressed in multiple OSN subtypes in addition to DII-projecting cell types, each of which give rise to distinct glomeruli in specific domains of the dorsal bulb, apparently breaking the class-based cell-type restriction regime. The innervation patterns observed in **Figures 2C-D** are remarkably similar to those of 5×21-OR1A1 axons in **Figure 1G**.

### Differentially expressed gene analysis

#### Topological framework for OR expression analysis

The approximately 10 million OSNs populating the mouse MOE express almost 1,100 intact OR genes (Saraiva et al., 2015). Expression of these ORs is not uniformly dispersed throughout the entire MOE, but rather is restricted to regions of the MOE, often referred to as zones. Several papers have ascribed numerical appellations to the zones (Miyamichi et al., 2005; Ressler et al., 1993; Vassar et al., 1993), using topological anatomical boundaries. An RNA-seq-based bioinformatic approach calculated a continuous zone index (1.05-5.00) along the dorsoventral (DV) axis, subdividing the MOE into 38 DV indices (DVIs) or “zones” (Tan and Xie, 2018) plus an “unusual location”, previously defined as the “patch,” based on the expression pattern of the Olfr155-Olfr159 genes (Strotmann et al., 1992). According to this nomenclature, Olfr151 is expressed in the most dorsal region of the MOE, with a likely DVI of 1.05. In our dataset, 1,040 ORs have been assigned to a specific DVI.

#### OR genes are differentially expressed in the 5×21-OR1A1 line

Recent RNA-seq studies in the mammalian WOM have demonstrated that this high-throughput technology is highly reliable in quantifying the abundance of thousands of OSN subtypes, based on the chemosensory receptor (e.g., OR, TAAR) they express (Ibarra-Soria et al., 2017; Ruiz Tejada Segura et al., 2022; Saraiva et al., 2015; Saraiva et al., 2019). In the 5×21-OR1A1 mouse line approximately 13% of all OSNs express OR1A1 (based on glomerular volume). If the MOE does not compensate for the overrepresentation of a single OR with the generation of additional OSNs, then we hypothesize that the expression of other OSN subtypes will be reduced commensurately.

To test this assumption, we started by performing RNA-seq on the MOE of hemizygous 5×21-OR1A1 and WT male mice. We processed three biological replicates per mouse line. On average, each sample yielded between 41.8-62.1M total reads, of which 80.2%-93.4% mapped uniquely to the mouse genome. We detected approximately 1,150 ORs in MOEs of both 5×21-OR1A1 and WT animals. To gain further insight into the transcriptomic changes associated with the over-representation of the 5×21-OR1A1 transgene, we performed differential expression analyses (**See Table 1**). Genes were classified as differentially expressed if they showed significant fold change (FC) greater than 2 (i.e., ABS(log_2_FC)>1.0, p-adjusted<0.05). Using this stringent criterion, we identified 425 differentially expressed genes (DEGs) in total, of which 77 were overrepresented and 348 genes are underrepresented in the 5×21-OR1A1 line. Of the 77 overrepresented DEGs, 40 were Class II OR genes while none were Class I ORs or TAARs. Of the 40 overrepresented Class II ORs, 33 were ventrally expressed (DVIs 2.00-3.60), four (including Olfr151) were dorsally expressed (DVI 1.05), one was in the unusual location index, and two were not assigned DVIs by Tan and Xie. Due to the design of the 5×21-OR1A1 transgene (homo sapiens OR CDS), which contains the 5’and 3’UTRs of the endogenous Olfr151 gene (129 mouse OR gene), the most significantly increased gene in the 5×21-OR1A1 animals was Olfr151 (log_2_FC=7.23 increase; pAdj 6.3×10^−22^), as anticipated (**Figure 3A**). We confirmed the DESeq findings by RT-qPCR analysis of RNA levels for the Class I OR, Olfr690 and the Class II OR, Olfr358, both of which showed a ∼4-fold decrease consistent with the ∼6-fold DESeq data (**Figure S1**). Interestingly, A second OR1A1 line (Omura, 2021) with significant dorsal expression produced with IRES-Gcamp6f also revealed the similar ∼4-fold decrease in expression for Olfr358. On the other hand, 325/348 DEGs with diminished representation are OR genes, with a subset representing all the known dorsal classes (6 TAARs, 90 Class I ORs, 179 Class II ORs|DVIs 1.05-1.90). 227 of the underrepresented ORs have a DVI of 1.05. In total, the analysis shows that ORs comprised the genes with the most significant FC in RNA representation in the MOE (**Figure 3A, B**). Comparison of the cumulative normalized mean count data for all dorsally indexed (DVIs 1.05-1.9) ORs shows an approximate 56.8% reduction in expression in 5×21-OR1A1 mice compared to WT animals. Additionally, we found that markers for immature OSNs are more abundant and mature OSNs are less abundant in 5×21-OR1A1 animals, suggesting that a more rapid turnover of OSNs is taking place in these animals; fewer axons with an underrepresented OR identity are sent to the bulb, resulting in unstable glomeruli and, as a result, increased OSN loss (Ebrahimi and Chess, 2000).

**Figure 3.**
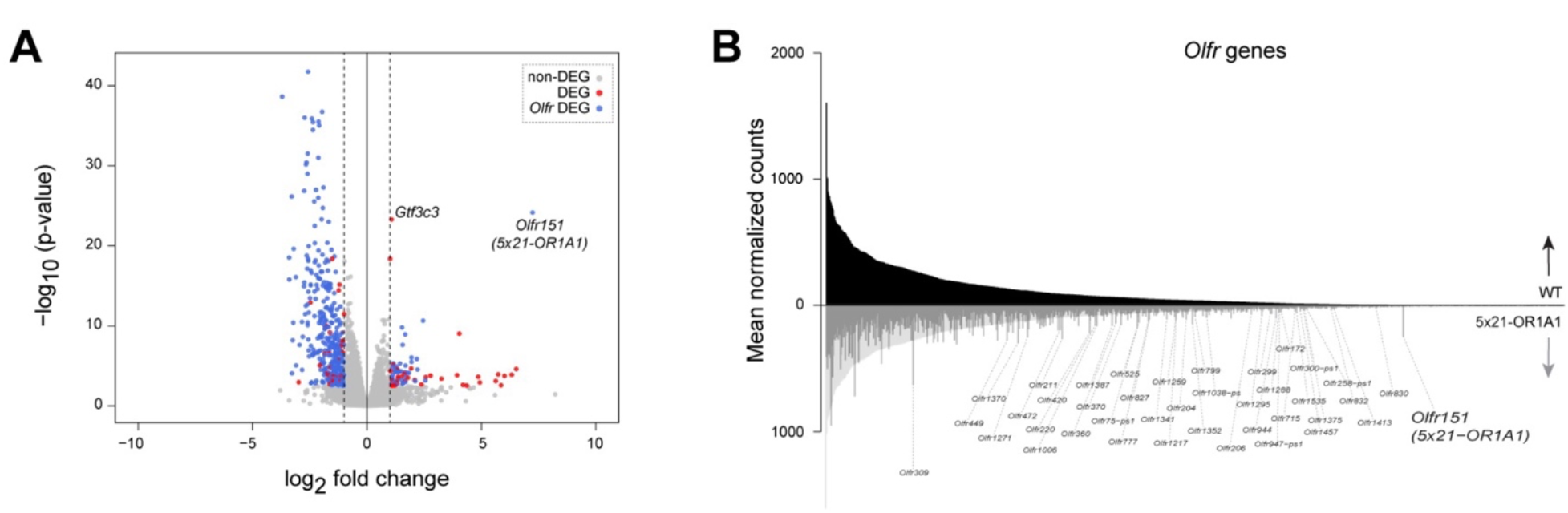
Differential expression analysis comparing the transcriptomes of 5×21-OR1A1 versus WT animals. (A) Volcano plot representing the -log10(p-value) versus the log2 FC for all expressed genes. Statistically significant changes (ABS(log_2_FC)>1.0, p-adjusted<0.05) in gene expression are marked by red-filled in circles for non-OR genes and by blue-filled circles for OR genes. Of the OR genes that met the statistically significant twofold change cutoff, the vast majority were underrepresented, whereas only a few OR genes were overrepresented. The Olfr151 gene was the most overrepresented transcript in our dataset due to the use of its UTR in the transgene backbone. One non-OR gene, Gtf3c3, which resides adjacent to the transgene in the 5×21-OR1A1 line’s genome, was significantly overrepresented as well (**Figure S2**). (B) Mirror Barplot for all OR genes. Mean normalized expression values of WT (top and bottom grey shading) and 5×21-OR1A1(bottom) are shown. OR genes are ordered by decreasing abundance based on WT. Most of the OR genes have some reduction of expression, except for the 40 labeled OR genes which have statistically significant twofold change overrepresentation.

Together, the RNA-seq data are consistent with our observations from wholemount imaging and support the hypothesis that the 13% of OSN representation allocated to the expression of the transgene OR1A1 is due to its ability to access the common OR choice mechanism in normal OSNs across cell types which are otherwise committed to the expression of other endogenous receptors.

#### TAARs are differentially expressed in the 5×21-OR1A1 line, but not Ms4a genes

TAAR genes are a third class of chemosensory receptors expressed in OSNs that primarily respond to amines and are situated in a single gene cluster on mouse chromosome 10 (mm10; chr10:23,920,406-24,109,582). All but two genes, Taar6 and Taar7b, are expressed in the dorsal MOE. TAARs are subject to singular gene choice like Class I and Class II ORs (Fei et al., 2021; Shah et al., 2021). As shown in **Figure 1B**, ΔTAAR4-YFP+ cells are located in the dorsal MOE and have a DVI of 1.05. Of note, Taar2, Taar7e, Taar7f, and Taar8b share this same DVI with Taar4 (Tan and Xie, 2018) and all five have approximately twofold underrepresentation in 5×21-OR1A1 mice compared to WT animals (**Figure 4A**). Ventrally expressed Taar6 (DVI 2.6 log_2_FC=1.15 decrease; pAdj 1.69×10^−02^), also shows underrepresentation in 5×21-OR1A1 mice. The remaining TAARs (Taar3, Taar7a, Taar7d, Taar9) are known to be dorsally restricted based on in situ hybridization (Pacifico et al., 2012), but were not captured in Tan and Xie’s analysis and thus have no assigned DVIs (Tan and Xie, 2018). Our DEG analysis shows that in total, 8 out of the 15 TAARs are significantly underrepresented (ABS(log_2_FC)> 1.0, p-adjusted<0.05) in 5×21-OR1A1 mice compared to WT animals. An additional five TAARs are underrepresented but have less than a twofold change (**Figure 4A**). Consistent with these results, when a ΔTAAR4-YFP line is crossed to the 5×21-OR1A1 line, we observe glomeruli on the dorsal bulb with smaller spherical radii when compared to βTAAR4-YFP-only animals, indicative of fewer coalescing axons of a TAAR identity (**Figure 4B**). The decrease in TAAR gene expression levels did not correlate to their location within the TAAR cluster. We also observed coexpression between ΔTAAR4-YFP and 5×21-OR1A1(Cherry) in 10/984 cells from 3 animals (∼1%) revealing that TAAR cell types can express the transgene (**Figure 4B – left inset)**. FISH probes for Taar2, Taar4, and Olfr690 (Class I) were rarely observed in 5×21-OR1A1 tissue whereas all three were readily observed in WT tissue (**Figure 4B – right inset**).

**Figure 4.**
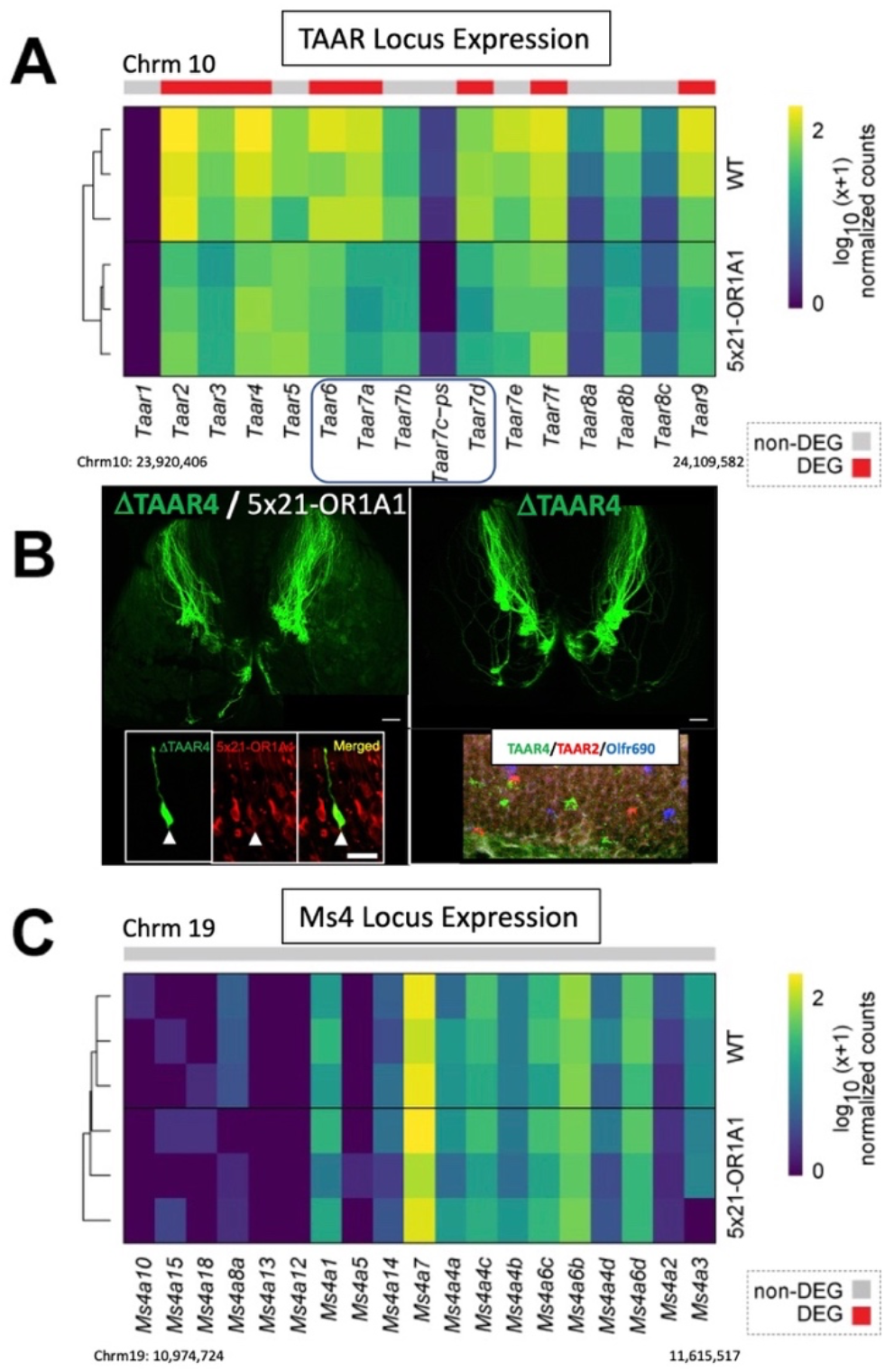
Expression profile of TAAR and Ms4a genes in WT and 5×21-OR1A1 samples. (A) Heatmap and phylogeny visualizing normalized expression of TAAR genes. RNA expression levels are represented on a log10scale of normalized counts (NC+1). Nearly all TAAR genes showed decreased representation. TAAR 2, 3, 4, 6, 7a, 7d, 7f, 9 are underrepresented two to five fold, padj < 0.05 (red labels above gene, see legend). Additionally, TAAR 7b, 7e, 8a, 8b, 8c also show reduced representation over WT. Blue box indicates TAAR genes expressed in ventral DVIs. (B) 5×21-OR1A1 crossed to ΔTAAR4-YFP (TAAR axons-green) reveals glomeruli with smaller spherical radii to ΔTAAR4-YFP-only animals, signifying fewer coalescing axons of a TAAR identity and consistent with the reduced expression of TAAR genes from DEG analysis. Left inset – Cross section of MOE reveals a ΔTAAR4-YFP OSN that is also positive for 5×21-OR1A1 (arrowhead). Right inset – RNAscope probes for TAAR2 (red), TAAR4 (green) and Olfr690 (blue) were readily observed in dorsal MOE of WT animals (n=5 sections) and almost impossible to find in 5×21-OR1A1 (n=10 sections). (C) Heatmap and phylogeny visualizing normalized expression of Ms4a genes in WT and 5×21-OR1A1 animals. None of the 19 genes show altered expression.

Recently, a non-GPCR family of chemosensory receptors which is not subject to singular gene choice, membrane-spanning, 4-pass A (Ms4a) genes (mm10; chr19: 10,974,724-11,615,517), was shown to express in necklace OSNs in the mouse MOE (Greer et al., 2016). In contrast to the TAARs, none of the 19 Ms4as met the statistical threshold for differential expression between 5×21-OR1A1 and WT animals (**Figure 4C**). These data further validate that the 5×21-OR1A1 transgene is being expressed through the mechanism of singular gene choice in chemosensory receptor expressing cells.

#### Differential expression of Class I ORs

Class I ORs are represented by 158 annotated OR genes and pseudogenes residing in a single cluster of approximately three megabases on Chromosome 7 in mouse (mm10; chr7:102,476,772-105,369,355), with the majority of Class I expressing OSNs constrained to the dorsal MOE. The Class I OR locus is split into two halves, separated by the hemoglobin beta chain complex (Hbb) that resides between Olfr67 and Olfr66 (**black arrow, Figure 5**). Tan and Xie assigned only 115 of 157 Class I ORs to DVIs due to 42 ORs with expression too low to be assigned. From their wild-type data, they assigned 110 of the Class I ORs to a DVI of 1.05 (Tan and Xie, 2018), which is consistent with prior studies showing Class I ORs to be dorsally restricted.

**Figure 5.**
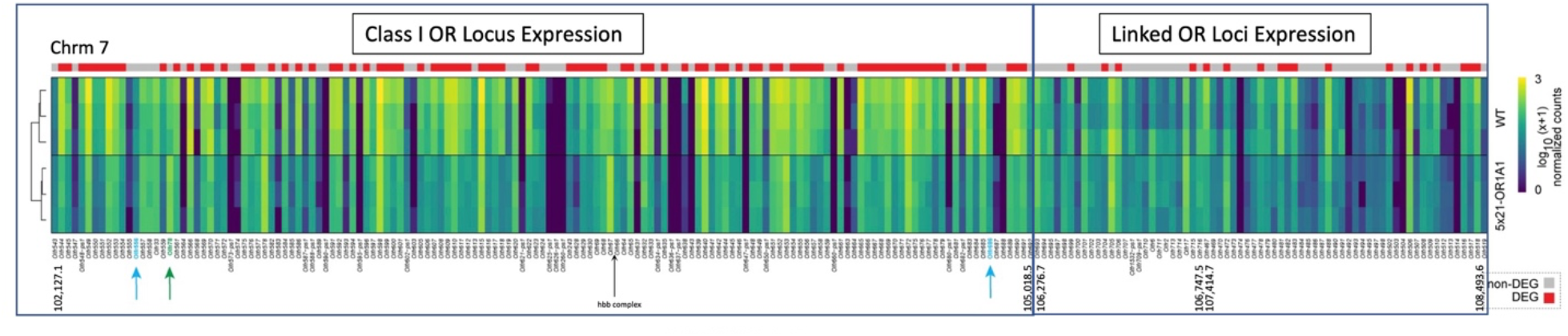
Heatmap and phylogeny visualizing normalized expression of genes in the Class I OR locus and two adjacent OR loci in WT and 5×21-OR1A1 animals. Nearly all expressed Class I OR genes (88/115) with assigned DVIs showed underrepresentation, padj < 0.05 (red labels, see legend); two of the Class I OR pseudogenes were among them. Only Olfr78, highlighted green, reveals modest upregulation (log_2_FC=0.71 increase; pAdj 9.31×10^−02^). The two non-classical Class I ORs, Olfr556 and Olfr686 expressed in Class II Cell types (blue labels) within DVI of 1.7 and 2.0 did not meet the cutoff stringency for differential expression. Only a few OR genes in the two linked OR clusters (1-2Mb downstream) are underrepresented, and nearly all of these ORs belong to DVI 1.05.

Our RNA-seq data mapped back to 145 of the genes in the Class I cluster and showed that Class I OR underrepresentation in 5×21-OR1A1 animals is equally distributed 1.5Mb centromeric and telomeric to Hbb. In 5×21-OR1A1 animals 87/110 DVI 1.05-expressed ORs showed significant underrepresentaion while only 1/3 non-DVI 1.05 Class I genes showed significant underrepresentation: Olfr543 (DVI 1.25 log_2_FC=0.41 increase; pAdj 8.09×10^−01^); Olfr550 (DVI 1.4 log_2_FC=2.70 decrease; pAdj 5.97×10^−10^) [less expressed]; Olfr571 (DVI 1.9 log_2_FC=0.07 decrease; pAdj 9.78×10^−01^) (**Figure 5**). Two other ORs identified as Class I based on coding sequence phylogeny, Olfr556 (DVI 1.7 log_2_FC=0.96 decrease; pAdj 2.49×10^−01^) and Olfr686 (DVI 2.0 log_2_FC=0.83 decrease; pAdj 8.17×10^−02^), were previously determined to be expressed in Class II OSNs (Bozza et al., 2009) and their expression levels do not reach the DEG threshold in 5×21-OR1A1 animals (**blue arrows, Figure 5**). Only one Class I OR, Olfr78, is modestly overrepresented (log_2_FC=0.71 increase; pAdj 9.31×10^−02^) but does not meet the fold change stringency to be classified as DEG-enriched (**green arrow, Figure 5**). Proximity to the Class I enhancer J/eA or the two putative OR enhancers-Greek Islands Milos and Pontikonisi is not correlated with any pattern of under or over representation for Class I ORs.

The wide-spread differential expression observed is not affecting the overall genomic region as the most proximally linked OR cluster, 750kb away (mm10; chr7:106,675,004-107,150,581), shows only 3/25 ventrally-expressed ORs (DVIs 2.4, 2.6, and 3.05) are underrepresented in 5×21-OR1A1 animals (**Figure 5**), whereas the next linked OR cluster, 2.4Mb away (chr7:107,812,262-108,894,420), has 13/42 underrepresented ORs with 11 belonging to DVI 1.05, similar to nearly all Class I ORs.

In comparison to Olfr78, the only non-Class I OR gene in DVI 1.05 to be DEG-enriched in 5×21-OR1A1 animals is Olfr420 (log_2_FC=1.53 increase; pAdj 1.97×10^−08^), which represents the second most overrepresented OR gene (based on significance) in our dataset. Interestingly, both overrepresented genes, Olfr78 (Class I) and Olfr420 (Class II) have been shown to have non-olfactory gene expression in sensory ganglia (Manteniotis et al., 2013). Also, like Olfr78, Olfr420 is flanked by two DVI 1.05 OR genes that are both significantly underrepresented in 5×21-OR1A1 animals, Olfr424 (log_2_FC=2.25 decrease; pAdj 3.17×10^−16^) and Olfr419 (log_2_FC=0.59 decrease; pAdj 8.84×10^−2^). Based on this, it appears that the proximal promoters of both Olfr78 and Olfr420 are unaffected by the presence of the 5×21 enhancer element.

#### Differential expression of Class I/II ORs relative to Dorsoventral index

Overall, our sequencing results show that 518 ORs are expressed in the entire dorsal MOE: 110 Class I ORs, 212 Class II ORs, and 5 TAARs (327 total) that retrospectively map to DVI 1.05, while 191 expressed ORs map to DVIs 1.25 to 1.9. Our data is consistent with ORs from three isogenic strains published by Ibarra-Soria and colleagues that we retrospectively mapped to DVIs: 314 ORs with DVI of 1.05 and 184 ORs with DVIs 1.25 to 1.9 (Ibarra-Soria et al., 2017). There are 496 ventrally expressed genes with a DVI 2.0 to 5.0 in our data set and 481 in the Ibarra-Soria data set. The 5×21-OR1A1 transgene produced underrepresentation of 275 ORs from the dorsal MOE comprising 8 DVIs: 227 OR genes from DVI 1.05 and 48 OR genes from DVIs 1.25-1.9. In contrast, only 45 OR genes for the remaining 31 DVIs (2.0-5.0 and unusual location) were underrepresented (**Figure 6**) (all DVIs as assigned by Tan and Xie). Taken together, two trends emerge from Tan, Ibarra-Soria, and our data. First, DVI 1.05 has the most ORs assigned to it. Second, DVIs from 1.25 to 5.0 each have fewer assigned ORs leading to a greater representation of OSNs expressing a particular gene in the more ventrally indexed OSNs, inferred by greater mean FPKM and assuming equal number of cells per index (**Figure S2**). Cell count data from 11 OR knock-in mice confirm this trend as the two most ventrally expressed OR KIs, Olfr124 [SR1] (DVI 4.3) and Olfr15 [MOR256-17] (DVI 3.05), have the most numbers of associated neurons (Bressel et al., 2016). Finally, Ibarra-Soria’s data showed that average FPKM counts for each OR from the three isogenic strains used reveal that DVIs 2.0-5.0 have 3.5x more counts compared to ORs with DVIs of 1.05-1.9, suggesting that only 22% of OSNs reside in the dorsal zone, with DVI 1.05 representing the majority of dorsally-expressed ORs at 14.4% (Ibarra-Soria et al., 2017).

**Figure 6.**
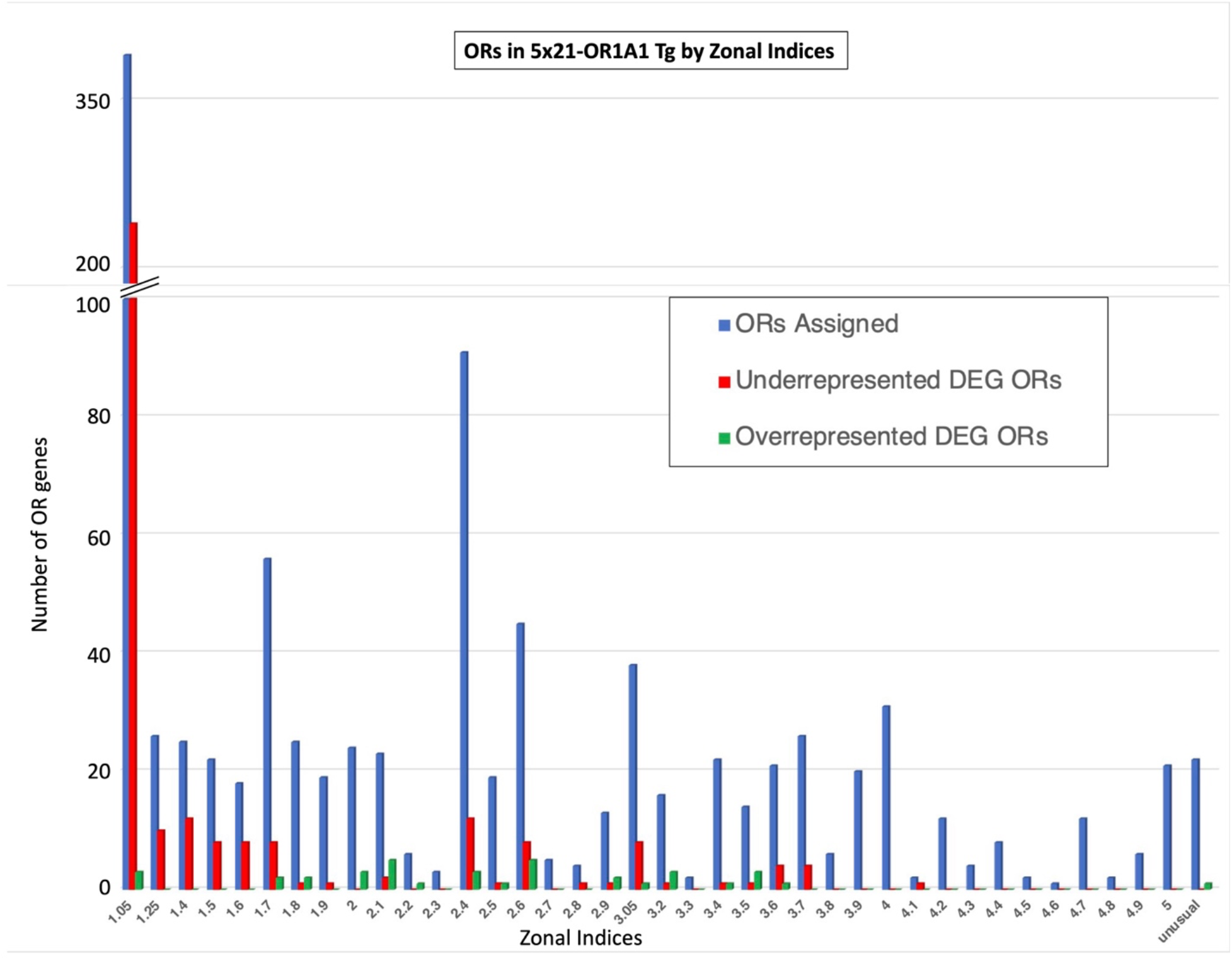
Altered representation of OR genes in 5×21-OR1A1 line by DVI. Counts of OR genes expressed in our data set by the 39 DVIs between 1.05-5.0 and the “unusual location” (Tan and Xie, 2018). Blue bars indicate the total number of OR genes assigned to the index. Red bars indicate the number of ORs within the DVI that are underrepresented and green bars indicate the number of ORs within the DVI that are overrepresented based on criterion for DEG (ABS(log_2_FC)> 1.0, p-adjusted<0.05). The 5×21-OR1A1 transgene has the greatest proportional effect on the ORs expressed in the indices of the dorsal MOE. Most of the underrepresented OR genes are from the dorsal DVIs between 1.05-1.7, wheras the 40 overrepresented OR genes are distributed between all DVIs. 392 OR genes (including TAARs) were not assigned DVIs by Tan due to low expression and are not reflected above. Excluding Olfr151, five of these OR genes were underrepresented and two were overrepresented in our data.

Conclusive evidence shows that the effect of preferential selection of the 5×21-OR1A1 transgene over Class II OR genes occurs throughout the olfactory subgenome and is not restricted to a single cluster or on a single chromosome (**Figure 7A**). As shown above, Class I ORs and TAARs (predominantly DVI 1.05 expressed), which reside in single clusters on different chromosomes, chromosomes 7 and 10, respectively (**Figure 7A, B**) are substantial underrepresented in OR1A1 mice. We mapped the remaining 139 underrepresented OR genes in DVI 1.05, which are Class II, and find them distributed among nearly all chromosomes and clusters (**Figure 7B**). The 44 OR genes in our data set from DVIs 2.0-3.9 that were underrepresented were not linked by chromosome or cluster either (**Figure 7B**). The most ventral aspect of the MOE, DVIs 4.0 to 5.0, contains 99 OR genes that were poorly represented in our data set; likely due to the difficulty in dissecting tissue. From this set of genes only one OR gene was significantly underrepresented.

**Figure 7.**
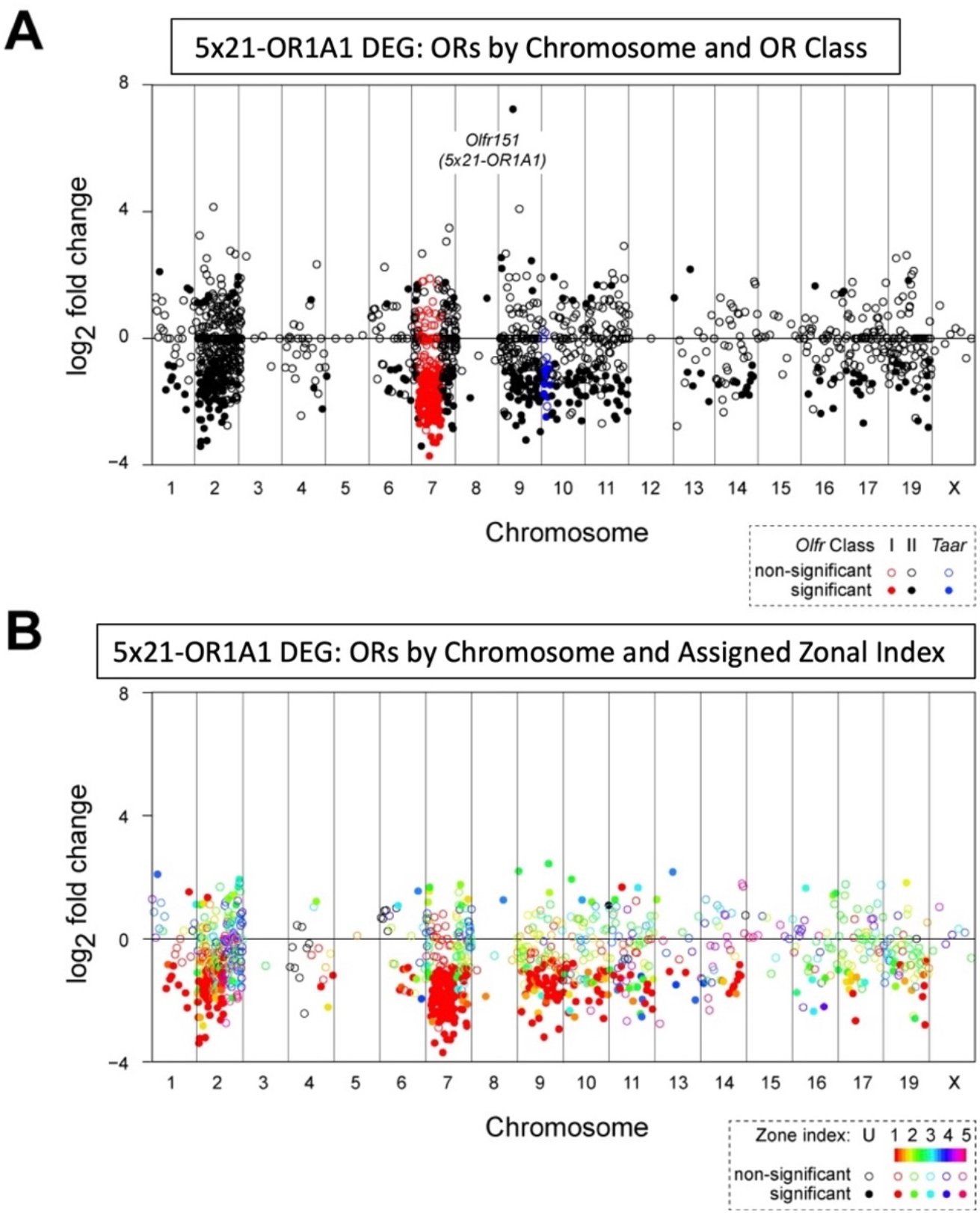
Scatter plots of differential expression analysis for all Class I, Class II, and TAAR genes expressed in the MOE comparing 5×21-OR1A1 to WT samples. Represented are the log_2_FC of expression for each gene in relation to its chromosome of origin and chromosomal position, from left to right (not to scale). (A) Class I OR genes (red) map to a single cluster on Chromosome 7. TAAR genes (blue) map to a single cluster on Chromosome 10. All other ORs genes are phylogenetically Class II (black) and map to nearly all chromosomes, with the exception of Chromosome 18 which contains no OSN-specific chemosensory receptor genes. Solid-colored circles represent differentially expressed genes (ABS(log_2_FC)> 1.0, p-adjusted<0.05). Class II DEG are distributed on all chromosomes and not linked to the DEGs observed for Class I and TAAR genes. (B) Scatterplot, same as in (A) with chemorecptor genes color-coded by DVI (Tan and Xie, 2018) and reveal that DVIs between 1.05-1.6 genes are the most affected across all chromosomes. None of the 392 unassigned ORs are shown, including Olfr151.

Both 5×21-OR1A1 and 4×21-Olfr151β transgenes utilize a strong gene choice enhancer (x21) to improve the probability of choice for the Olfr151 promoter. Olfr151 is poorly expressed in 129 mice and is pseudogenized in B6 mice. As a result, Olfr151 is not assigned a DVI by Tan and Xie. Olfr160 [M72] is 96% identical to Olfr151 but is expressed in all mouse strains and is assigned to DVI 1.05. Knockin data provides support that Olfr151 exhibits the same zonal patterning as Olfr160 (Vassalli et al., 2002). Our data show that the x21 sequence has amplified the DVI 1.05 probability to an extent where it not only increases the representation of the Class II OR transgene in DVI 1.05 Class II cells, but also gains probability of choice in Class I and TAAR cells in DVI 1.05. Additionally, the x21 element showed the ability to expand its Class II OR probability into Class II cells in DVIs of the MOE ventral to DVI 1.05.

#### DEGs as a function of the Greek Islands

The 5×21 sequence has been derived from the core HD-site of the H element, with homology to a 19-mer sequence found in the P element enhancer. The H element is known to associate with a set of sequences known as Greek Islands. The Greek Island sequences have been proposed as a mechanism for the nucleation of expression of one Class II OR allele per OSN through trans-acting aggregation of an enhancer hub (Markenscoff-Papadimitriou et al., 2014; Monahan et al., 2017). In this model, the 5×21 sequence would require the Greek Island hub to activate its expression in Class II OSNs in order to be expressed in a singular manner.

In contrast, the Greek Island hub is not associated with Class I or TAAR gene expression. However, our 5×21-OR1A1 DEG data shows these genes to be robustly underrepresented (62% Class I and 38% TAAR), implying that the Class II transgene has been chosen for singular expression in Class I and TAAR cells. We also observe the formation of glomeruli in the DI and DIII domains of the OB in 5×21-OR1A1 animals. This indicates that the 5×21 element not only has a Class II character but is also able to gain probability of choice in all known non-Class II OSNs without having to go through the Greek Island hub (Cichy et al., 2019). If the 5×21 element does not require the Greek Island hub to gain probability of choice in Class I and TAAR cells, then it stands to reason that a common mechanism for monoallelic expression exists which underlies choice in all OSN classes. As a result, the Greek Island hub may not be mechanistically relevant to choice and may only be an incidental aggregation of OR regulatory elements within the Class II OR compartment.

#### DEG of non-OR genes and proximity to the 5×21-OR1A1 transgene

Only two non-OR genes, Musc5Ac and Cyp2c44, are found to be underrepresented within the set of the first 100 differentially expressed genes in 5×21-OR1A1 animals, revealing that the specificity of x21 action is restricted to OR gene expression.

Tan and Xie also assigned DVIs to 712 non-OR mouse genes (a reduced subset of 202 genes was produced when they applied greater stringency). It should be noted that Tan and Xie classify TAAR genes as non-OR genes while we chose to include them as a class of chemosensory GPCR genes. Only 3/22 underrepresented non-OR genes in our data set are assigned to a DVI in the Tan lower stringency data set: Nqo1 (DVI 1.05 log_2_FC=1.20 decrease; pAdj 1.98×10^−13^); Ivl (DVI 1.4 log_2_FC=1.18 decrease; pAdj 8.58×10^−03^); Cyp2c44 (DVI 1.4 log_2_FC=2.06 decrease; pAdj 3.97×10^−04^). In 5×21-OR1A1 animals, expression of Nqo1, a known marker of mature dorsal OSNs, is decreased 2.5-fold, consistent with either loss of Class I neurons and/or increased turnover. Looking at the 37 non-OR genes that were upregulated, only two were assigned DVIs: 5430403N17Rik (DVI 2.6 log_2_FC=1.14 increase; pAdj 3.43×10^−04^) and Gm26794 (DVI 3.05 log_2_FC=1.79 increase; pAdj 8.21×10^−03^).

The 5×21-OR1A1 transgene was tandemly integrated within the Hecw2 gene on Chromosome 1 between ∼49,170,017-53,904,772 removing 4Mb of DNA. If our transgene contains a probability of choice enhancer, then perhaps it could have increased expression for one or several of the genes in close proximity to the site of integration. The integration of 5×21-OR1A1 removed all proximal genes upstream of Hecw2, leaving a gap of 3Mb until the next cluster of genes. The two genes downstream of Hecw2 are Ccdc150 and Gtf3c3 at 54,478,185 (**Figure S3**). The greatest upregulated gene in our analysis was Olfr151, followed by Gtf3c3 as the second most probable upregulated gene (log2FC=1.07 increase; pAdj 4.23×10^−21^) in the genome (based on significance) (**Figure 3A**). Gtf3c3 is normally expressed at very low levels in globose basal cells and in the OMP layer of the MOE as shown by RNA hybridization. In 5×21-OR1A1 animals, expression of Gtf3c3 could be observed in 5×21-OR1A1 OSNs and globose basal cells (**Figure S4**). These data strongly suggest that the multimerized x21 enhancer increases probability of expression for linked genes within OSNs. When such a linked gene is an OR or a surrogate OR, such as β2AR, then axon guidance can ensue forming glomeruli (Pyrski et al., 2001).

## Discussion

### Expression patterns of ORs in the olfactory epithelium

Expression of the phylogenetically distinct Class I OR genes is restricted to OSNs in the most dorsal index of the MOE, DVI 1.05, with the exception of three Class I OR genes in the set of those assigned DVIs: (Olfr543 (DVI 1.25), Olfr550 [MOR16-1] (DVI 1.4), Olfr571[MOR21-1] (DVI 1.9)). Class I OSNs also group their axonal projections to the most dorsal regions of the OB, domain I (DI)..

Class II expressing cells are found across all 39 DVIs, unlike Class I expressing OSNs which are predominantly restricted to DVI 1.05. However, we still see restricted expression of Class II ORs to specific DVIs. Given the broad distribution but zonal restriction of expression of Class II ORs within Tan’s 39 DVIs between 1.05 to 5 throughout the MOE, it is reasonable to hypothesize that there are multiple types of Class II OSNs. We anticipate that each of these sub-Class II cell types will establish their own probability of choice for a subset of Class II receptors that will have the opportunity to be expressed within the sub-class. We think that it is likely that these sub-Class II cell types are in some way defined by the DVI. Evidence that suggested the existence of multiple sub-Class II cell types was previously shown using the Olfr6 [M50]→Olfr17 [P2] coding sequence swaps and Olfr151 transgenes (D’Hulst et al., 2016; Feinstein and Mombaerts, 2004).

We show that 5×21-OR1A1 reduces the expression of Class I, II and TAAR genes and is capable of projecting axons to each of the dorsal domains in the OB (DI, DII, and DIII), as defined by the formation of distinct glomeruli in each of the domains, suggesting that the transgene is chosen in all classes of OR and TAAR-expressing cells (Figure 1). In addition, we observe that multiple distinct glomeruli form within the DI and DII domains in the OB. Because axons with like OR-identity projecting from the same cell-type should coalesce into a single, homogeneous glomerulus within its given class domain in the OB, we propose that the nine distinct glomeruli we observe with OR1A1 identity, two Class I and seven Class II, are in fact the result of OR1A1 axonal projections from several distinct cell types. This also implies that because the multiple glomeruli are situated within the DI and DII domains of the bulb, there are likely multiple sub-classes of Class I and Class II which coincide with the multiple cell types observed and are best identified by their topological DVI. In addition, we show the formation of three distinct glomeruli of 4×21-Olfr151β identity within the DI, DII, and DIII domains in the OB by axonal projections from OSNs confined to the dorsal MOE (**Figure 2**). Taken together, we propose that this is indicative of the ability for both transgenes to break class restriction and gain probability of choice and expression in each of the three main classes of OSNs.

### Zone or Cell Type?

What defines the epithelial expression domain of an OR? Is the proximal promoter of a given OR deterministic of expression within distinct cell types (Class I, II or TAAR) or do they produce unequal probabilities of choice in different cell types that only allow OSNs which chose them to be sustained in a specific zone of the MOE? For example, are the Olfr556 (DVI 1.7) and Olfr686 (DVI 2.0) genes regulated by “Class II promoters” or do they have certain probabilities of being expressed in both Class I and Class II cell types, but the Class I probability is insufficient to sustain their expression in Class I cells in DVI 1.05? All phylogenetically similar Class I ORs are linked in a 3Mb cluster and are regulated by the J/eA enhancer, except for Olfr556 and Olfr686, which are not differentially regulated in J/eA knock outs (Cichy et al., 2019). One could postulate that the Class II expression of those two genes could be regulated by a Class II enhancer, like the Greek Island Pontikonisi, which contains both core Lhx2 binding site motifs (TAATGA and TAATTG) and is located 2.2 kb downstream of Olfr556, similar to other Class II control elements. Alternatively, it may be that Olfr556 and Olfr686 have the potential to be expressed in Class I cells, however their proximal promoters have such a low probability for expression in cells within DVI 1.05 that there are never enough OSNs with those identities in DVI 1.05 to form stable dorsal glomeruli to sustain them. Our new conceptual framework, as demonstrated by the formation of multiple homogeneous glomeruli by OSNs expressing our high-probability transgenes, 5×21-OR1A1 and 4×21-Olfr151β, and DEG analysis of 5×21-OR1A1 animals versus WT, suggests ORs may have certain probabilities of expression between different cell types that are modulated by cell-type restriction factors and are only sustained in a given region if they achieve the critical mass necessary to form stable glomeruli, but are able to tap into a common mechanism of singular gene choice that is present for all OR-expressing OSNs.

### The H element enhancer is governed by the x21 sequence element and can tap into the common mechanism for singular gene choice

It was previously shown that the 2kb H element, derived from the cis-enhancer of the ventrally expressed OR genes, can impose increased dorsal representation for the Class II Olfr16 [MOR23] (DVI 1.05) transgene, However, the projection site of the resultant glomeruli appeared to skew to a large glomerulus in the most dorsal aspect of the OB where the Class I domain (DI) resides as well as to a smaller set of axonal projections to the endogenous Olfr16 glomerulus in DII (Vassalli et al., 2011). Like the full H-element, we hypothesize that the multimerization of the 21bp core HD sequence produces a cooperative effect that increases the probability that a driven transgene can access the singular gene choice mechanism common to all OSNs.

The core 2kb H element contains 3xTAATGA homeodomain-type recognition sequences in a span of 92 bases. A small knockout of 187bp that included all three HD sites abolished functionality of the element in mice (Nishizumi et al., 2007). Previously we observed that a sequence with homology (13/16 identities) to one of these HDs within the P element could increase the probability of choice for a linked gene (Vassalli et al., 2011). Our x21 sequence is not artificial and contains an extended HD homology to the 19mer derived from the P element. It has previous been proposed that synergistic affects between HD and O/E sites lead to increased OR choice, yet our x21 sequences is devoid of an O/E-like site. Genomic analysis of the olfactory sub-genome has also found x21-like sequences in both Class I J/eA, and TAAR TE1/TE2 enhancers, consistent with the idea of a shared mechanism of gene choice (Cichy et al., 2019; Fei et al., 2021; Iwata et al., 2017; Shah et al., 2021).

### Cell type restriction of ORs

Based on our data, we conclude that OR class (Class I, Class II, or TAAR) does not define cell type, but the correlation of the class expressed within a given cell type is unlikely to be coincidence either. Cell type restriction is a mechanism that seems likely to group OR functionality to the same projection sites in the bulb (**Figure S5**). In this regard, grouping odor receptive properties of ORs may be a mechanism to more accurately characterize the nature of a stimulus, consistent with the fact that the spatial location of ORs correlate with sorption patterns of the odorants they recognize (Ruiz Tejada Segura et al., 2022). Therefore, cell type restriction would not necessarily be a means to reduce the repertoire of OR genes an OSN can chose from in order to reduce the complexity governing the process of singular gene choice. Rather, cell types would allow neighboring glomeruli in close proximity in the OB to more accurately compare similar odors.

### Probability of choice of an OR promoter

By comparing the effects of the Olfr151 transgene with and without the high probability x21 enhancer, it appears that OR promoters have an inherent probability of choice within a cell type, which also correlates to an expression region in the MOE corresponding to a DVI. Our data suggest that as probability of choice rises for a given OR promoter, so does its ability to be selected by additional cell types. For example, a dorsally-expressed gene would naturally have a “lower” probability to be chosen in a more ventral region owing to the “dorsal character” of its promoter elements. However, if its probability for choice is increased, it gains the ability to be chosen in ventral cell types by tapping into the mechanism of gene choice common to all OSNs. If enough OSNs of this identity project axons to form ectopic glomeruli, these cells will be sustained and the choice can be observed. It is currently unknown what drives these cell-type restrictions and correlated zonal expression patterns, but it seems likely to be controlled by regulatory sequences.

The profound change in expression of Class I and TAAR genes by the x21 driven transgene is not the result of competition with Greek Island enhancers as they have not been implicated in the regulation of OR genes in those cell types. One Class I gene, Olfr78 seemed to be refractory to the effects of x21 and showed a modest upregulation, suggesting that its probability went up when Class I competition for expression went down. This OR gene likely has a special set of enhancers (Zhang et al., 2016) that helps to maintain its expression. For Class II genes that are said to be regulated by Greek Islands in cis, we do not observe protection from overrepresentation of the x21 sequence, suggesting that the x21 enhancer has higher probability of choice than cis-acting Greek Islands.

Previous experiments have used the tTA-TetO synthetic promoter system to drive high levels of OR expression in the MOE that phenocopies elements of olfactory receptor gene choice in an attempt to unlock the mysteries of singular gene expression of ORs in OSNs (Abdus-Saboor et al., 2016; Fleischmann et al., 2008; Nguyen et al., 2007). The limitations of these experiments are due to the onset and efficacy of tTA activation, which shows similarities to but does not access the mechanism of singular gene expression. The difference between the expanded representation of our high probability transgenes in the MOE and the overexpression seen in the synthetic tTA-TetO expression system is that our transgenes tap into the inherent mechanism of probability of choice that exists in the endogenous OSN. Cells choosing our transgenic 4×21-Olfr151β allele produce similar levels of mRNA and protein as OSNs choosing the endogenous Olfr151 allele and coalesce axons into the same glomeruli. In contrast, there is no evidence that supports tTA-driven TetO-Olfr151 is expressed at endongenous levels; in fact, it has been shown that their axons do not coalesce with endogenous Olfr151 axons into the same glomeruli, implying that the axons poses different identities (Ma et al., 2014). The x21 singular gene choice element, derived from the H element, relies on the cell’s natural expression of Lhx2. Our transgenes represent an additional allele to the over 2,000 OR alleles in the genome available to be chosen by an OSN. This is not the case with the tTA expression system, where there is no competition within the genome for the binding of tTA to other regulatory elements; it will bind exclusively to TetO leading to the preferential expression of the gene under its control.

Furthermore, the tTA expression system is not subject to class and zonal restriction regimes associated with OR gene choice as it sits outside the singular choice mechanism. Using OMP or CAMKII-tTA drivers after the onset of endogenous OR expression to express TetO-driven ORs can supplant the expression of the endogenously chosen OR allele and leads to the TetO-driven OR’s expression throughout the MOE. This “bully” phenomenon is commonly referred to as post-selection refinement (Abdus-Saboor et al., 2016). The PSR-phenotype produces thousands of regularly spaced glomeruli with axons of dual identities; that of the originally expressed OR, which influences axon guidance, combined with the supplanting TetO-driven OR. By contrast, our x21 enhancer is expressed at the onset of endogenous OR expression, through the mechanism of singular gene choice, and is not causing the formation of thousands of small, dual identity glomeruli. Rather, the multiple glomeruli observed in x21 animals within the DI and DII OB domains are very large with a unique identity derived from high overrepresentation of OSNs singularly expressing the transgene in DVI 1.05. In addition, expression of the x21-driven transgene in ventral DVIs of the MOE, resulting from increase probability of choice, yield a string of solitary glomeruli in the ventral OB. Therefore, we contented that while tTA-TetO is a useful tool to study aspects of olfactory singular gene expression, it is insufficient to understand the totality of the mechanism, as it does not rely on the endogenous mechanism in OSNs for its expression, as do our high-probability enhancers.

## Conclusion

At lower probabilities of choice, ORs may be transiently expressed in cell types other than the one that it is normally found. OSN identity arises from both the expressed OR and some yet to be determined cell-type specific modifier giving rise to formation of unique glomeruli. If transient OR expression occurs outsides of its cognate cell type, then their axons must coalesce in sufficient numbers to form stable glomeruli, otherwise these cells will wither and die. It is in this way that cell-type restriction, as mediated by probability of choice, for a certain repertoire of ORs creates the appearance of stochastic expression to a particular DVI. The mechanisms of glomerular formation provides an additional level of cell-type selection. The value of Tan and Xie’s Zonal Indices is likely that these 39 demarcations (DVIs between 1.05-5.0 plus the “unusual location”) correlate to OR peak probabilities of expression and may indeed reveal the total number of cell types (**Figure S5**).

Through the use of high probability elements, our data provide a new conceptual framework to view OR gene choice as the probability of an OR being expressed in a given cell type, which in wild-type animals lies within a domain of expression in the MOE. Our high probability x21 enhancer appears capable of breaking the cell-type restriction regime, allowing it to be chosen for stochastic expression in a broad range of cell types throughout the MOE leading to multiple distinct glomeruli and revealing a common mechanism of gene choice that transcends OR classes.

## Supporting information

Table 1_DEG_Analysis

DEG_Raw Data

## Acknowledgments

We thank the Hunter College Animal Facility Manager Barbara Wolin and Veterinarian Patricia Glennon for help in maintaining the transgenic colony, and Rada Norinsky at the Transgenic Core Facility at The Rockefeller University for generating transgenic founders. We thank WCMC genomic core facility for sequencing. Many thanks to Zach Gershon technical support in RNA extraction. Many thanks to Cami John-Melendez for artwork in supplemental Figure S5.

## Data and materials availability

Mice available at Jax: **STOCK Tg(Olfr151-OR1A1**,**-MAPT/mCherry)D736Feins/J;** Stock No: **029600**

## Material and Methods

Wholemount analysis 4 to 12 weeks old mice were used for the analysis by wholemount confocal microscopy imaging with LSM510 (Zeiss). Whole-Mount Immunohistochemistry Fix the sample in 1% Paraformaldehyde (PFA) at 4°C on a shaker overnight. Rinse the sample with fresh 1x Phosphate Buffered Saline (PBS) at least three times to remove excess PFA. Block the sample in blocking solution (4% Bovine Serum Albumin, 0.5-2% Triton-X 100, 5% Normalized Goat Serum, 1x PBS) at 4°C on a shaker overnight. Add the sample to the primary antibody solution (1% Bovine Serum Albumin, 0.5% Triton-X 100, 5% Normalized Goat Serum, 1x PBS, 1:1000 Primary Antibody) and wash at 4°C on a shaker overnight. Wash the sample with fresh 1x PBS at least three times at 4°C on a shaker for approximately two hours each time to remove excess primary antibody. Add the sample to the secondary antibody solution (1% Bovine Serum Albumin, 0.5% Triton-X 100, 5% Normalized Goat Serum, 1x PBS, 1:400 Secondary Antibody) and wash at 4°C on a shaker overnight. Wash the sample with fresh 1x PBS at least three times at 4°C on a shaker for approximately two hours each time to remove excess secondary antibody.

### RNA extraction and sequencing

Dissected MOE processed with miRNeasy Mini Kit (50) # 217004, QIAGEN. mRNA was prepared for sequencing using the TruSeq® RNA Library Prep Kit ((NonStranded and Poly-A selection, Illumina). All samples were sequenced on an Illumina machine, to generate single-end 100 bp sequencing reads and had an average depth of 49.23 ±3.00 (SEM) million reads. Newly generated RNA-seq data from these studies is available in the GEO dataset under the study accession number XXX

### Bioinformatics analysis

The reads qualities were assessed for each sample using FastQC. Sample Fastq files were aligned to the mouse reference genome Mus Musculus GRCm38.p5 (GCA_000001635.7) using TopHat2 (v 2.1.1) with 2 mismatches. Reads were retained if uniquely mapped to the genome, multi-mapped reads were removed from further analysis. We used HTSeq-count (0.9.1, -t exon and –m union) to obtain the number of reads that were mapped to each gene in the Gencode M16.

Bigwig files were generated from bam files for visualization using bam2wig.py (RSeQC package v3.0.1), all the bigwigs were normalized by Reads per Million (RPM) to compare across samples of different library size. We used Bioconductor package – DESeq2 (v1.22.2) to import raw HTSeq counts for each sample into R statistical software (v3.5.3) to create a raw count matrix.

Sample Transcriptome data quality was assessed by evaluating homogeneity and similarity between samples. We used variance stabilizing transformation(vst) and regularized log transformation(rlog) functions (DESeq2 package (v1.22.2)) to transform the raw count data, (dist) function to calculate sample-to -sample distances. We plotted heatmaps of distance matrix along with PCA of 1000 most variable/most expressed genes to identify putative outlier samples. These samples were excluded from the following differential expression analyses. We also used DESeq2 (v1.22.2) to normalize the raw count matrix according to sequencing depth and RNA composition (median of ratios method), then to perform differential expression analysis comparing WT vs 5×21-OR1A1 samples.

### RNAscope^®^

Fluorescent in situ hybridization was performed on fresh frozen sections of 12 μm coronal cryosections mounted onto SuperFrost slides. Tissue was fixed in 4% PFA at 4ÅãC for 15 minutes. Slides underwent a dehydration series from of 50%, 70%, and 100% ethanol (v/v in water) for 5 minutes each. Several modifications to the RNAscope. protocol were made to optimize signal on sensitive tissue of the MOE. Slides were then pretreated using Protease IV diluted 1:30 in 1X PBS for 30 minutes and then rinsed 3 times in 1X PBS. Sections were then stained for specific mRNA targets using RNAscope. Protocol (Advanced Cell Diagnostics, RNAscope. Multiplex Fluorescent Assay Version 1 Reagent Kit For Fresh Frozen Tissues User Manual 320295-QKG). Probes were hybridized at 40C overnight in the HybEZ oven (Advanced Cell Diagnostics). After amplifier steps and washes sections were stained with DAPI for 30 seconds and then mounted with ProLong Gold Antifade and coverslips which were fixed to slides with nail polish. Slides were imaged using the Perkin Elmer UltraView spinning disk confocal microscope and viewed under 63X oil immersion. The target probes used in this assay are custom reagents designed by Advanced Cell Diagnostics and are currently available in the ACD catalog. Mouse target probes are as follows: 318431-C1 RNAscope^®^ Probe- Mm-Taar4-C1 853051-C2 RNAscope^®^ Probe- Mm-Taar2-C2 853061-C3 RNAscope^®^ Probe- Mm-Olfr690-C3 853041-C1 RNAscope^®^ Probe- Mm-Gtf3c3-C1 313791-C2 RNAscope^®^ Probe- Mm-Omp-C2 550241-C3 RNAscope^®^ Probe - Mm-Olfr151-upstream-C3

### Gene Expression Workflow

We dissected the olfactory epithelium from 6-week-old mice and stored the tissue at -80°C in RNA*later* Solution (Invitrogen, Cat#: AM7020). We extracted RNA with the QIAGEN miRNeasy mini kit (Cat#: 217004), a BEAD BUG6 homogenizer (Benchmark Scientific), and on-column DNase treatment (QIAGEN, Cat#: 79254). To make cDNA, we used 2.5 μg, SuperScript IV VILO, and an ezDNAse treatment step to definitively remove any DNA contamination (Invitrogen, Cat#: 11766050). Primer were designed using the Primer-BLAST tool on the NCBI website and relative quantification of cDNA was evaluated using SYBR-Green (Roche, Cat#: 04707516001) and a Roche LightCycler 480 Instrument II. Each sample/gene pair was loaded in triplicate and each experimental performed thrice. Log2Fold Change values were calculated with a modified ΔΔCt method using *Acsm4, Acss2, Slc25a35* as reference primers and corrected primer efficiencies. Additional information and the MiQE checklist are available in supplemental information or upon request.

### RT-qPCR Statistics and Gene Expression Reproducibility

To correct for multiple T tests, we used the Holm-Bonferroni method. The plating design, Log 2 Fold Change calculations, and statistical tests are available in supplementary files. R version 4.1.0 was used to write code to process raw Ct values into L2FC values, perform statistical tests, and generate graphs. Code can be made available upon reasonable request.

## Figure Supplement

**Figure S1:**
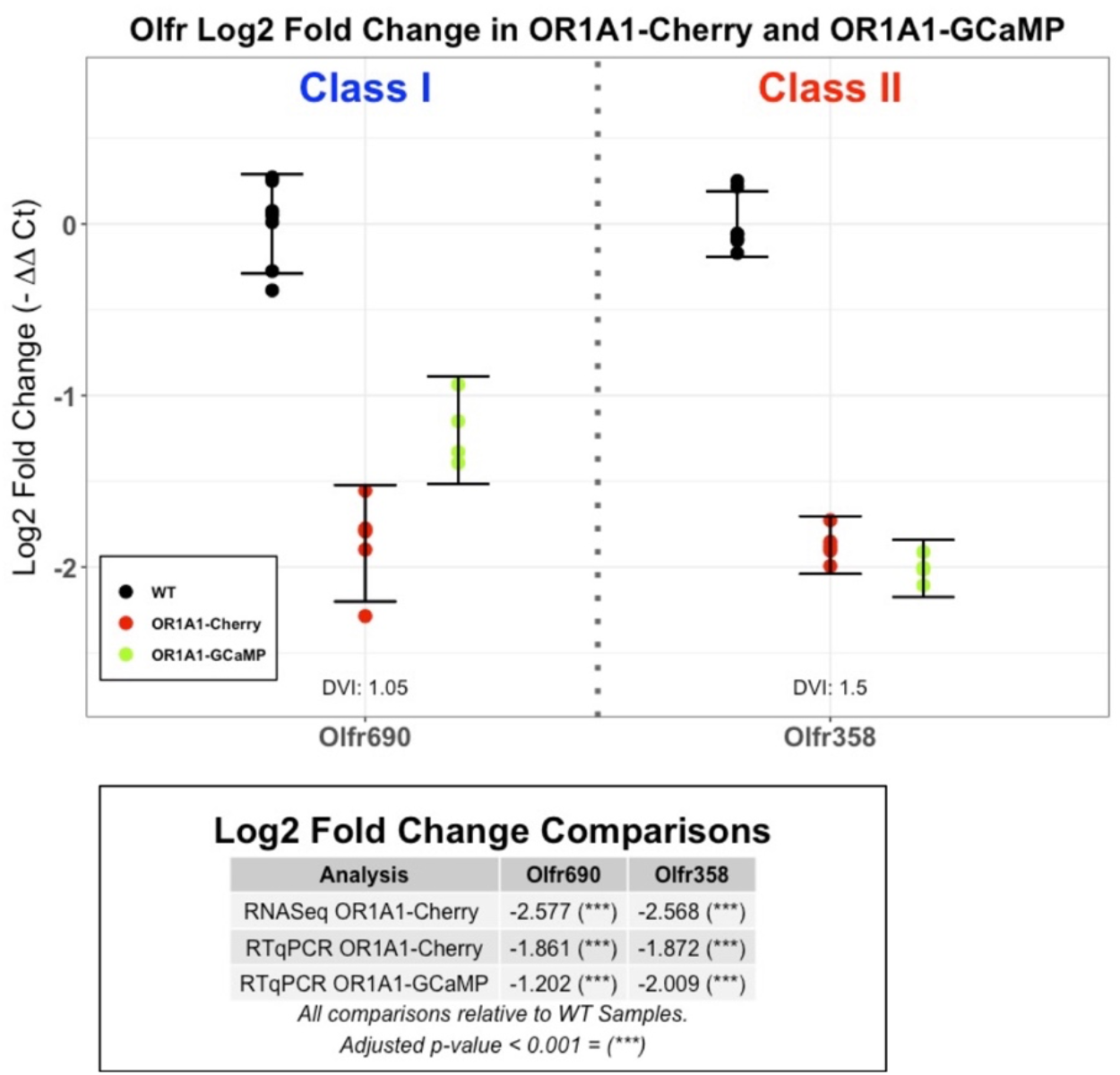
RT-qPCR for Class I and Class II Olfr genes in two OR1A1 lines. Log2 Fold Change (L2FC) of Olfr mRNA in two OR1A1 lines (5×21-OR1A1-IRES-tauCherry and 9×21-OR1A1-IRES-MP-GCamP6f (Omura, 2021)) relative to WT. L2FC was calculated using a modified ΔΔCt method with three reference genes (Acsm4, Acss2, Slc25a35). Two Olfr primer pairs were tested, with the Olfr Class and Dorsal/Ventral Index (DVI) displayed. Error bars indicate 95% confidence intervals for each mean. Biological replicates: WT N = 7, OR1A1-Cherry N = 5, OR1A1-GCaMP N =4. WT animals come from all lines; other genotypes come from their specific lines. Technical replicates: 3 experiments in triplicate. Alpha = 0.05. RT-qPCR of a subset of Olfr genes in OR1A1-Cherry is consistent with the results of DESeq in the same line. The RT-qPCR Log2 Fold Change values are relative to WT values. RT-qPCR p-values were corrected using the Holm-Bonferroni method. Alpha = 0.05.

**Figure S2:**
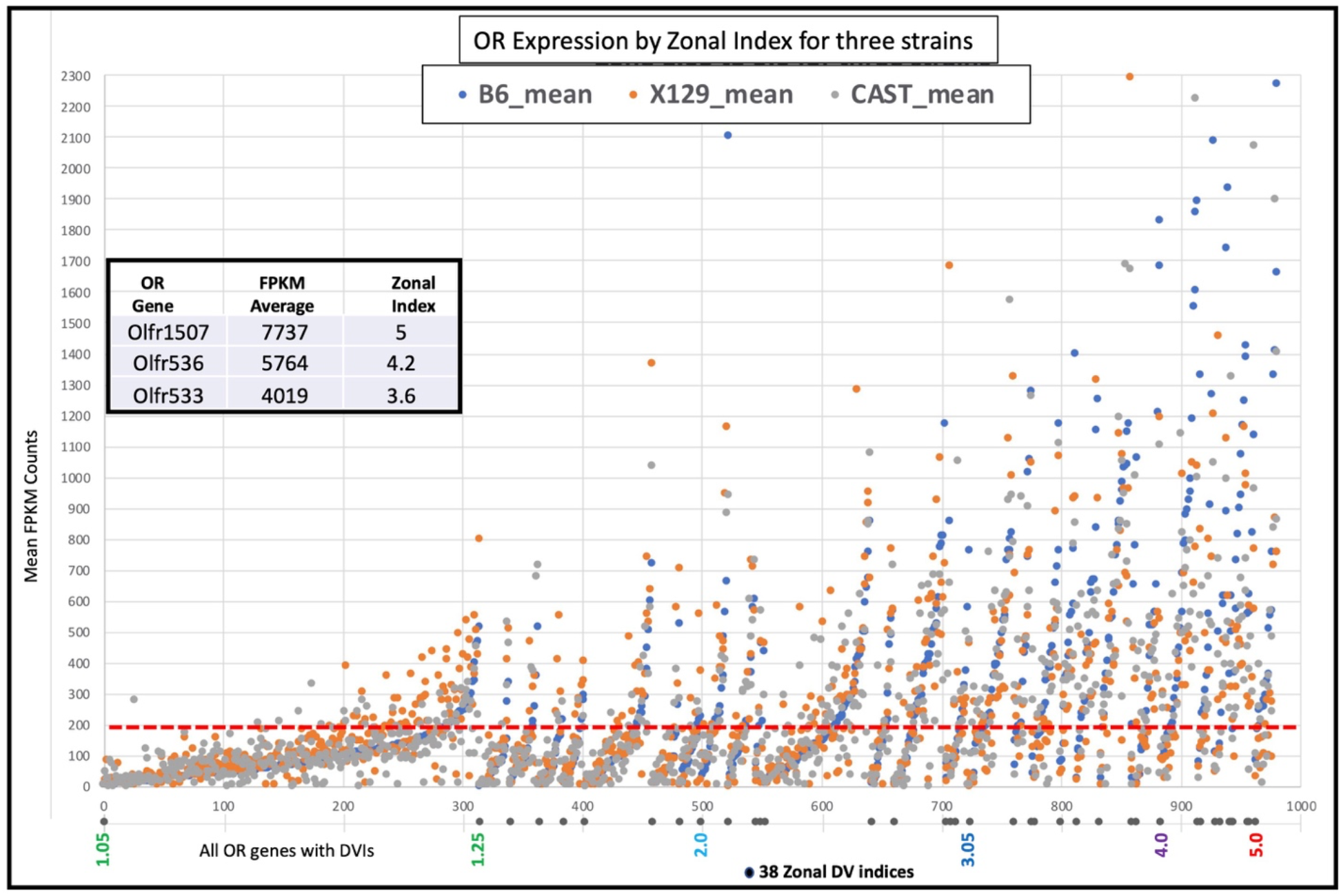
Distribution of mean FPKM expression levels for Class I and Class II ORs across three strains of mice: C57bl6, X129 and CAST. Scatter plot is organized by DVI and anchored by mean expression of C57bl6 within a Zone Index. From this set of 982 expressed genes, 314 are from a DVI of 1.05 and represent the lowest per OR expression within a zone. By contrast a DVI of 5.0 contains only 20 genes and represent the highest expression levels per OR. There is a trend from DVIs between 1.05 to 5.0 where fewer OR genes are assigned within progressively ventral DVIs, but each OR shows more representation. As an example, below the dotted red line at the 200 FPKM boundary, 92% OR genes (290/314) from DVI 1.05 exist compared to 15% (3/20) from DVI 5.0. Three OR genes have been left off of this plot (see inset) because all three strains show FPKM values above 2300: Olfr1507 (avg. FPKM: 7737), Olfr536 (avg. FPKM 5764) and Olfr533 (avg. FPKM 4019). Olfr1507, Olfr536 and Olfr533 are common between the top 4 represented OR genes from the three strains and expressed in the most ventral regions of the MOE.

**Figure S3:**
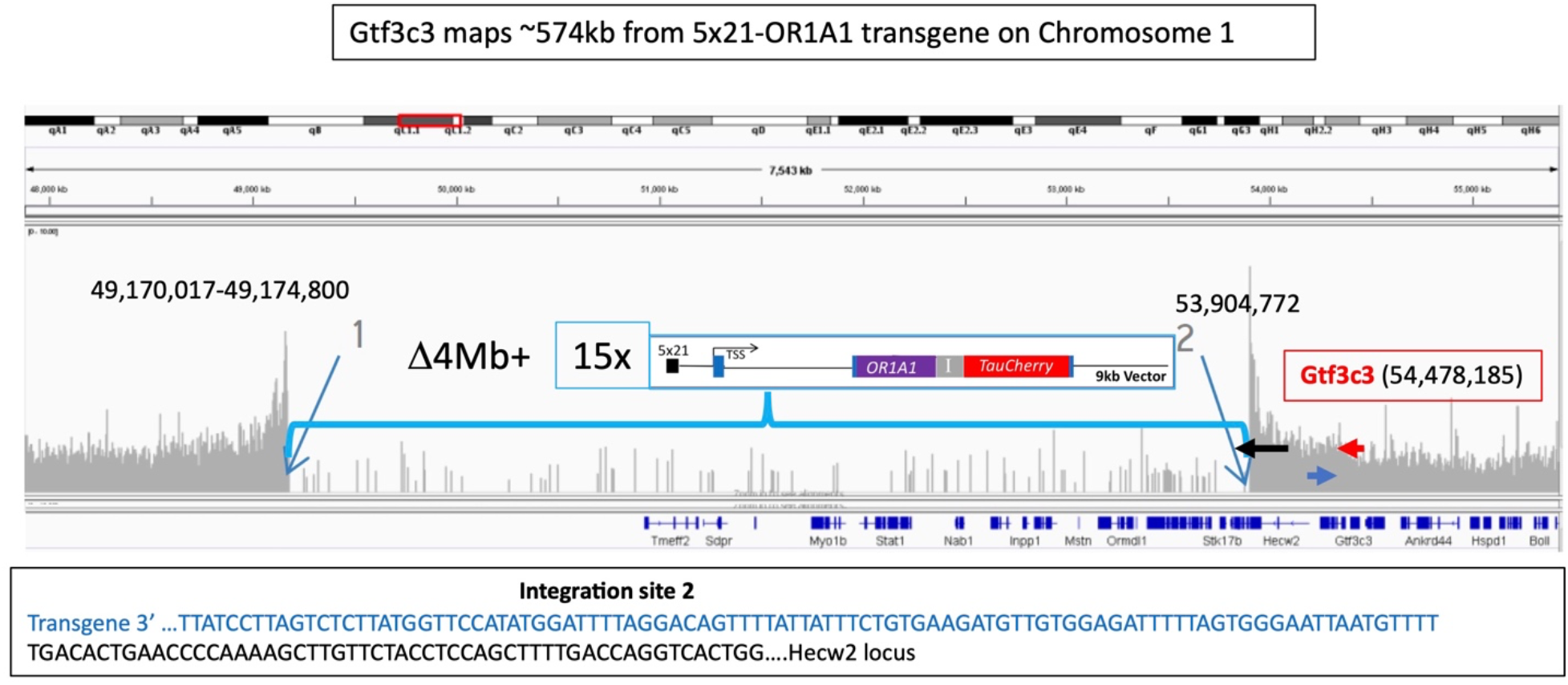
Chormosomal integration of the 5×21-OR1A1 transgene. TLA sequencing reads (Cergentis) map 15 copies of the 5×21-OR1A1 transgene to Chromosome 1 within the Hecw2 gene and were determined to reside in the 4Mb region deleted from ∼49,174,800 to 53,904772.. The copy number of the transgene derived by qPCR is 18, consistent with TLA mapping (D’Hulst et al., 2016). The Gtf3c3 transcription start site resides ∼584kb downstream of the transgene

**Figure S4.**
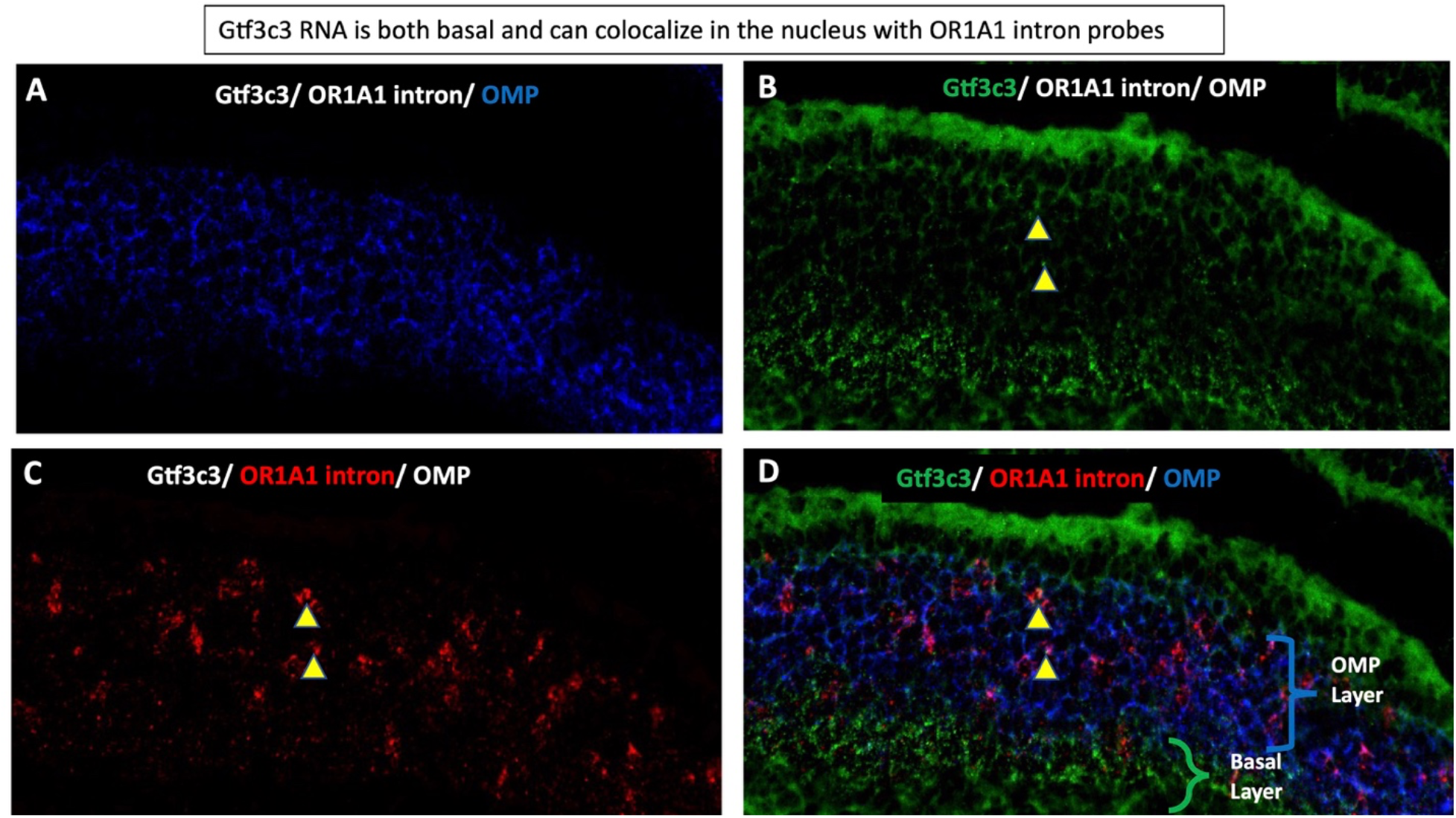
RNAscope analysis of Gtf3c3 in 5×21-OR1A1 animals. (A) OMP expression (blue) shows all mature olfactory neurons. (B) Gtf3c3 (green) expression reveals punctate expression in basal region and with low levels of expression in the OSN layer. (C). OR1A1 intron probe reveals nuclear region within OSNs where OR1A1 is being actively transcribed. (D) Overlay shows several OR1A1 expressing OSNs with Gtf3c3 expression in close proximity (yellow arrowheads).

**Figure S5.**
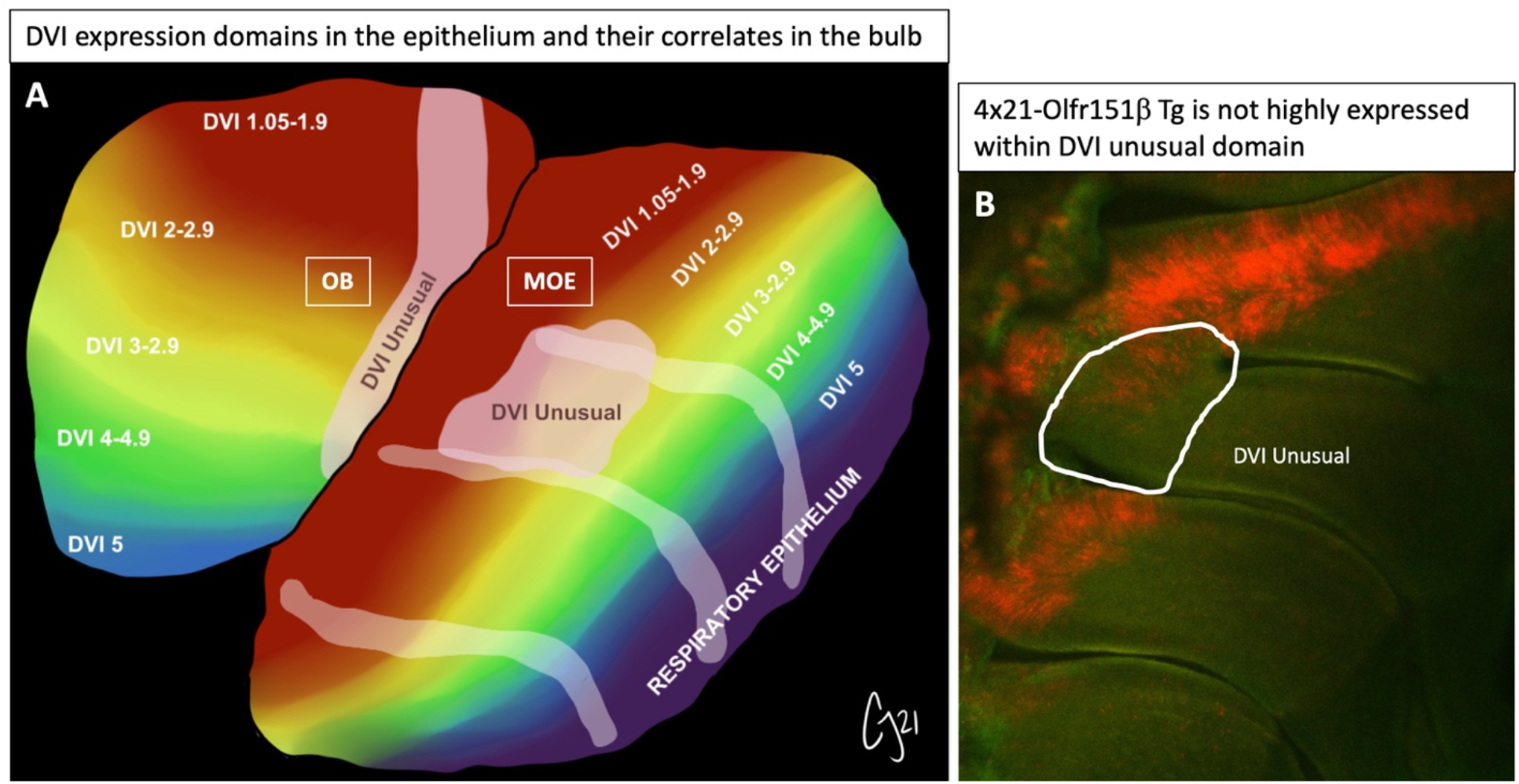
Schematic representation of DVIs as they correlate to gross projections to the olfactory bulb. (A). Dorsally expressed Class I ORs comingle in the MOE with Dorsal Class II ORs and the TAAR genes, but their projections segregate into distinct domains of the dorsal bulb. Of note, DVI unusual overlaps multiple DVIs from 1.05 to 3.05. (B) 4×21-Olfr151β transgene is expressed in the dorsal MOE but shows a marked reduction in expression within the DVI unusual domain.

